# The immunological profile of RC17 hESC-derived dopaminergic neural progenitor cells *in vitro*: implications for the STEM-PD clinical trial

**DOI:** 10.1101/2024.01.23.576826

**Authors:** Annabel J Curle, Shaline V Fazal, Shamma Qarin, Sarah K Howlett, Xiaoling He, Roger A Barker, Joanne L Jones

**Affiliations:** Clinical Neurosciences Department, University of Cambridge, Cambridge, CB2 0AH, England; Wellcome-MRC Cambridge Stem Cell Institute

**Keywords:** Parkinson’s Disease, Cell Therapy, Stem Cells, Human Embryonic Stem Cells, Immunogenicity, Clinical Trials, Regenerative Medicine

## Abstract

Parkinson’s Disease involves the progressive loss of dopaminergic neurons (DAn), prompting clinical trials replacing cell loss with neural grafts. This includes the transplantation of pluripotent stem cell-derived DAn progenitor cells (NPC) currently under investigation in the STEM-PD trial. To determine the likelihood of immune rejection post-grafting, we characterised the immunogenicity of the STEM-PD product (RC17-hESC-derived NPCs), comparing them to human foetal ventral mesencephalic tissue (hfVM) previously tested in trials, including our own TRANSEURO trial. Despite MHC-Class I expression, upregulated by proinflammatory cytokines, no immune response to NPCs was detected *in vitro*. Instead, they were immunosuppressive. Transcriptomic analysis revealed similarities between RC17-NPCs and hfVM, both strongly upregulating antigen processing and presentation pathways in response to IFNγ. Furthermore, immunosuppressant mycophenolate mofetil detrimentally affected NPC survival and differentiation *in vitro*. Overall, our data suggest that aggressive immunosuppression is not required following hESC-NPC transplantation and that caution should be exercised when selecting the immunosuppressive regimen.

## Introduction

Parkinson’s disease (PD) is a progressive neurodegenerative disorder characterised by the loss of A9 dopaminergic neurons (DAn) in the substantia nigra pars compacta (SNpc). Alongside non-motor symptoms, dopamine cell loss is responsible for significant motor deficits in PD, including bradykinesia and rigidity. Until now, managing these motor symptoms has involved the use of dopaminergic drugs such as levodopa, dopamine agonists and inhibitors targeting enzymes that break down dopamine. While usually highly effective in the early stages of disease, over time these drugs can cause their own set of side effects. These include neuropsychiatric and autonomic complications, and levodopa induced dyskinesias (LID) which may be disabling, necessitating more invasive interventions like deep brain stimulation (DBS). Consequently, there is a pressing need for more targeted, physiological replacement of dopamine.

One such approach is to graft new dopamine cells into the site of greatest dopamine loss within the striatum. Since the 1980s, different cell sources have been tested preclinically and in clinical trials. Of these, greatest success has been seen following transplantation of human foetal ventral mesencephalon (hfVM) tissue^1–24^. This tissue contains developing nigral dopamine cells of the type lost in PD. However, trial outcomes have been variable, likely due to differences in trial design, patient selection, amount of tissue grafted, and the level and duration of immunosuppression used^1,25,26^. Nevertheless, hfVM transplants have provided proof of principle that dopamine cell replacement therapy can be effective in PD^27,28^, with several patients showing marked clinical improvements with normalisation of dopamine levels in the striatum, patients coming off all their PD medication, and with evidence of dopamine cell survival up to 24 years post grafting^29^. Due to the limited availability of hfVM tissue, ethical issues associated with its use, and the failure to standardise such a therapy, the development of human pluripotent stem cell (hPSC)-derived dopamine cell products has been actively pursued^30–37^. Several induced pluripotent stem cell (iPSC)- and human embryonic stem cell (hESC)-derived dopamine neuron progenitor cells (NPCs) have now been demonstrated to work reproducibly well in animal models of PD^25,30,36–46^ with a good safety profile, which has now led to first in human trials^47–49^. The most recent of these is the European STEM-PD clinical trial initiated in February 2023 testing human ventral midbrain dopamine NPCs derived from the RC17 hESC line using the Nolbrant *et al.* protocol^32^ in eight people with Parkinson’s between Lund, Sweden and Cambridge, UK (NCT05635409^45^). This protocol enables the generation of NPCs in 16 days *in vitro*, at which point cells are cryopreserved until their time of implantation into patients.

One concern with any allogeneic transplantation is the immunogenic potential of the cells and subsequent possibility of immune rejection. Here we performed detailed immune characterisation of the STEM-PD product (RC17 hESC-derived NPC) comparing it to the previously successfully engrafted dissociated hfVM, tested in our previous TRANSEURO trial^50^. We studied the RC17 hESC-derived NPC *in vitro* at different stages of differentiation; including the undifferentiated hESC (day 0) to mature terminally differentiated DAn (day 45+). No NPC immunogenicity was detected *in vitro.* Instead, the NPCs had immunosuppressive properties reducing T-cell activation and proliferation.

Despite these reassuring findings, immunosuppressive drugs are, and are likely to continue to be, given post-transplantation. Therefore, we tested the effects of these drugs on NPC survival and differentiation *in vitro* ; this is rarely considered. We show that of the panel of drugs initially intended for the STEM-PD clinical trial, mycophenolate mofetil (MMF) negatively impacted the survival and differentiation of RC17-hESC derived NPCs into mature DAn. As a consequence, MMF has been removed from the STEM-PD immunosuppressive regimen.

Taken together our data suggest that grafted NPCs are at minimal risk of immune rejection, and that caution should be taken when selecting the immunosuppressive regimen of such studies.

## Results

### Generation of the STEM-PD product (RC17 hESC-derived NPCs)

First we confirmed successful differentiation of RC17 hESC to NPCs using the Nolbrant protocol^32^, as used in the STEM-PD trial. After 16 days of differentiation, RC17-NPCs were triple-positive for lineage commitment markers FOXA2, LMX1A, OTX2 as assessed by immunofluorescence (Sup Figure 1A) and flow cytometric analysis confirmed high expression of FOXA2, IAP and EN1 and a lack of PAX6 and OCT3/4 (Sup Figure 1C-D), confirming correct cell identity. A quantitative-PCR gene panel was used to further validate the expression profile of the day 16 NPC product, including those reflecting DAn progenitor state (*EN1, FOXA2, LMX1A, LMX1B, OTX1, OTX2*), ventral midbrain patterning (*PAX6, FOXG1, HOXA2, BARHL1*), positive predictors of dopamine differentiation (*CNPY1, PAX8, SPRY1, ETV5*), and mature DAn (*TH, NURR1, PITX3*), and a reduced expression of pluripotency genes (*OCT3/4, NANOG*) and negative predictor *FEZF1* (Sup Figure 1B). In all cases, gene and protein expression was comparable to, or higher than, that of the hfVM tissue. After differentiation for 45+ days to mature DAn, cells expressed the appropriate markers of differentiation including TH, TUJ1, FOXA2, and GIRK2 as detected by immunofluorescence (Sup Figure 1E) and were triple positive for TH, MAP2 and NURR1 as measured by flow cytometry (Sup Figure 1F-G). All of this was in line with the cell product being used in the STEM-PD clinical trial^45^.

**Figure 1.**
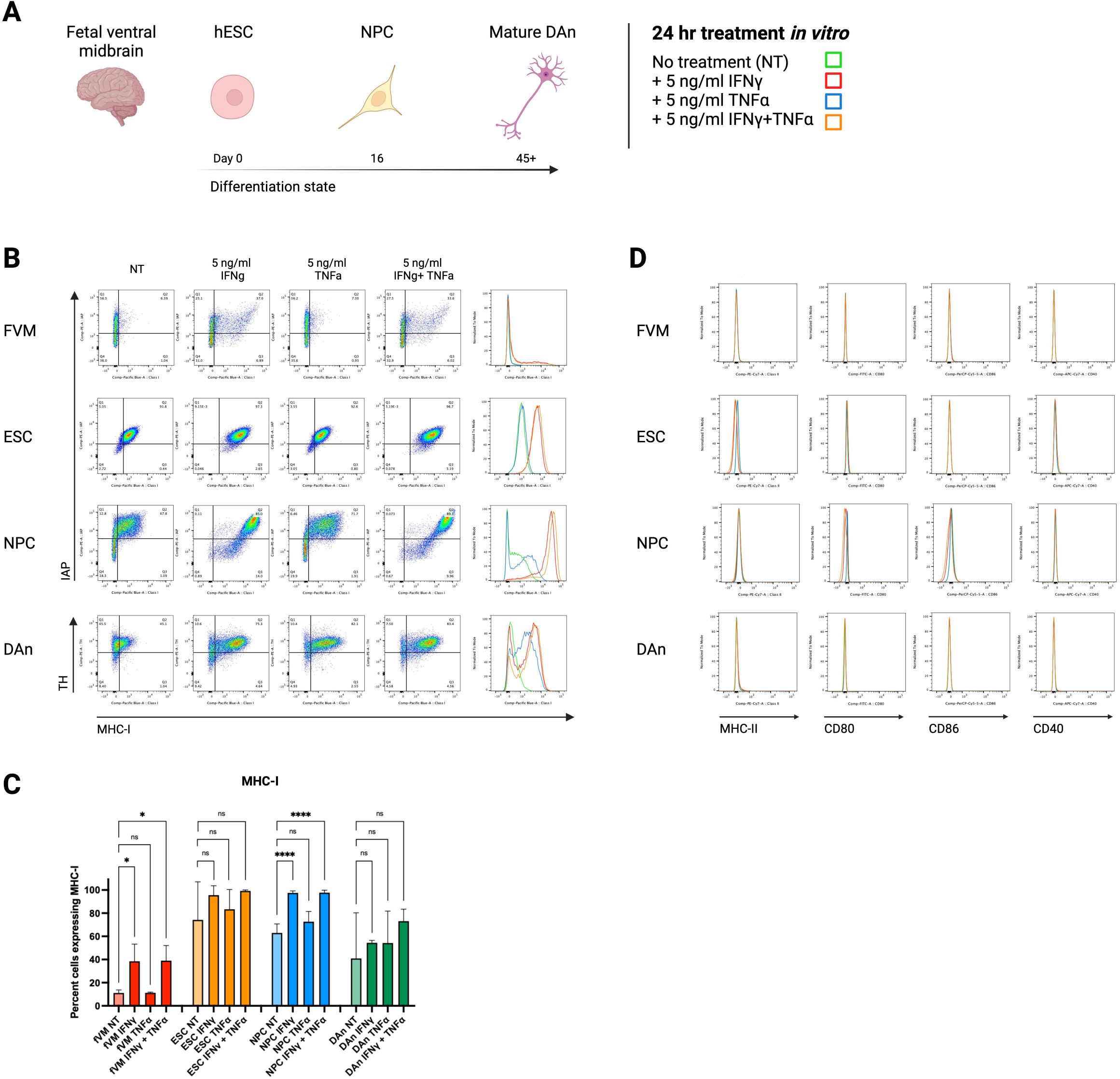
hESC-derived NPCs express MHC class I, which is increased by exposure to IFNγ. (A) hfVM, ESC, NPC, and DAn were seeded on appropriate coating media and left untreated (NT; green) or treated with 5 ng/ml IFNγ (red), 5 ng/ml TNFα (blue) or both IFNγ + TNFα (orange) for 24 hours before staining for surface protein expression. (B) Representative expression of MHC-I against cell type marker (y axis) and summary histogram overlays. (C) Quantification of percent cell expressing MHC-I with statistical comparison within cell type. (D) Histogram overlays of expression of MHC-II, CD80, CD86 and CD40 (D). Data are represented as mean ± STDEV. One-way ANOVA; n.s. = not significant *p<0.05, **p<0.01, ***p<0.005, ****p<0.001.

### hESC-derived NPCs express MHC class I, which is increased by exposure to IFNγ

Next, we examined the immunogenic potential of the NPCs by assessing their surface expression of immune relevant molecules by flow cytometry. To mirror the STEM-PD trial, the NPCs were assessed following freeze thawing and their profile was compared to dissociated hfVM tissue recently successfully transplanted in our completed TRANSEURO trial^50^. To determine if their immunogenic profile varies over time, undifferentiated hESC and day 45+ DAn were also examined. To mimic a likely, post-surgery, local inflammatory response and the known inflamed PD brain^51,52^, expression was also assessed after 24 hour exposure to the proinflammatory cytokines IFNγ and TNFα (Figure 1A, Sup Figure 2A) and IL-1β (Sup Figure 2A).

**Figure 2.**
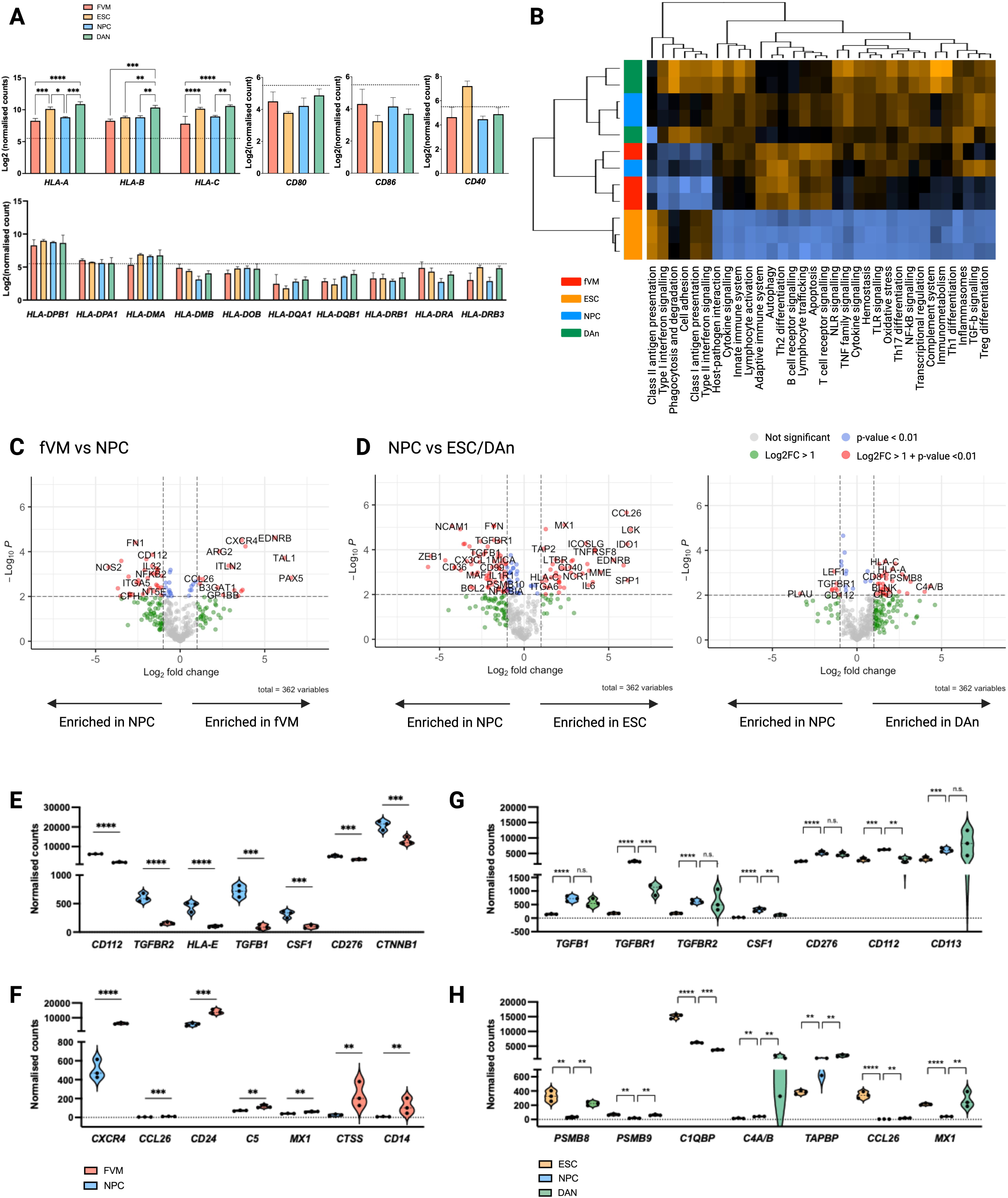
hfVM, NPCs and DAn are transcriptomically similar for immune relevant genes in non-inflammatory conditions. Nanostring nCounter analysis comparing hfVM, hESC, NPC and DAn. (A) Validation of Figure 1 surface expression at the transcript level; Log2(normalised count) quantification of MHC-I genes (*HLA-A, HLA-B, HLA-C*), MHC-II genes (gene names shown on x axis) and co-stimulatory genes *CD80, CD86* and *CD40*. (B) Clustered heatmap displaying immune-related pathways expressed by cells without treatment. (C-D) Differential expression plots comparing NPCs to hfVM (C), hESC and DAn (D) without treatment. (E-F) Violin plots depicting genes expressed at a significantly higher (E) or lower (F) level by NPCs than hfVM. (G-I) Violin plots depicting significantly differentially expressed genes by NPC than ESCs and/or DAn. Data are represented as mean ± STDEV. One way ANOVA; n.s. = not significant *p<0.05, **p<0.01, ***p<0.005, ****p<0.001.

The flow cytometric data displayed in Figure 1B shows MHC Class-I expression against the cell typing markers - integrin associated protein (IAP) for hfVM, hESC, and NPC and tyrosine hydroxylase (TH) for DAn. The summary data (Figure 1C) show percent MHC Class-I expression of the entire product (gated on live single cells) as would be transplanted into the patient. MHC Class-I was expressed by ESC, NPCs and DAn, with the percent expression higher than that seen in cells derived from the hfVM tissue (Figure 1B-C). 24-hour treatment (time and concentration course shown in Sup Figure 2) with IFNγ induced significant upregulation of MHC-I on hfVM and NPCs, and a small, non-statistically significant, increase on hESC and DAn. Treatment with TNFα alone had little effect (Figure 1B-C), and no clear additive effect when given in combination with IFNγ. Very little MHC-II expression was observed on any cell type. No upregulation was seen following exposure to IFNγ, TNFα or IFNγ + TNFα. No cell type displayed expression of the co-stimulatory molecules CD80, CD86 or CD40 (Figure 1D).

These data suggest that the NPCs may be more immunogenic than hfVM, particularly in an inflammatory environment, based upon their capacity to interact directly with CD8+ T-cells via their higher MHC-I expression. However, at no differentiation stage do they express all molecules required for direct T-cell activation.

### hfVM, NPCs and DAn are transcriptomically similar for immune relevant genes in non-inflammatory conditions

Having identified key surface molecule expression, we next performed targeted transcriptomic analysis on all cell types using the Nanostring nCounter Human Immunology V2 panel (which assesses the expression of 579 immunologically relevant genes) plus an additional 20 gene codeset designed to capture additional genes related to T-cell activation/inhibition and graft rejection (see pathway annotations and custom codeset in Sup Figure 3A). Cells were examined after 24 hours in culture, either with no treatment (NT) or following exposure to 5 ng/ml IFNγ for 24 hours - selected based on a preliminary experiment that, in line with the surface expression data, revealed stronger immune-related responses to IFNγ treatment compared to TNFα, and that combined treatment did not appear to be additive (Sup Figure 3B).

**Figure 3.**
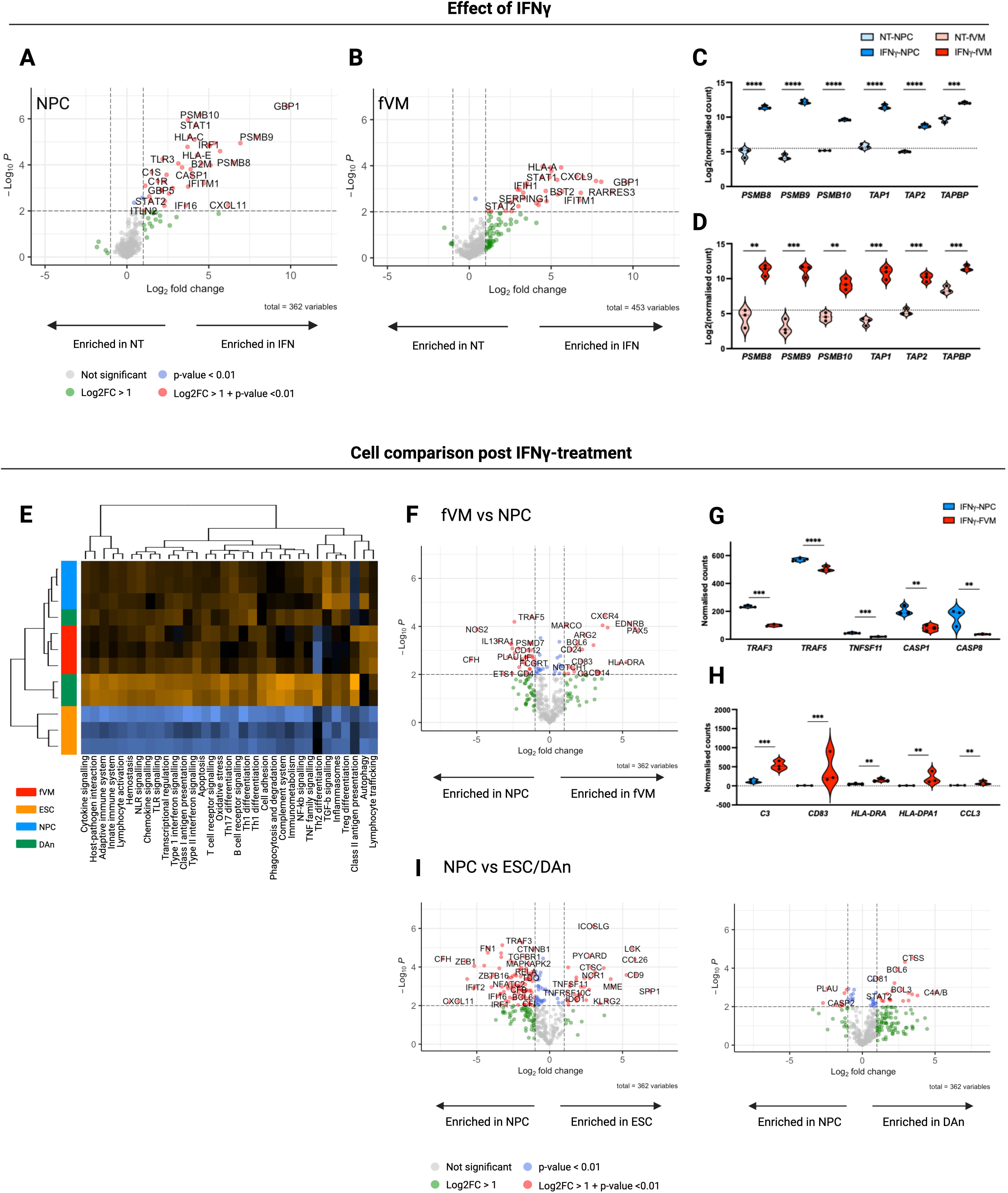
hfVM and NPCs both increased expression of antigen processing and presentationgenes in response to IFNγ exposure. (A-C) Effect of IFNγ on NPC and hfVM. Differential expression plot comparing (A) NPC and (B) hfVM stimulated with 5 ng/ml IFNγ for 24 hours against baseline of untreated cell type. (C) Antigen processing and presentation genes significantly upregulated by (C) NPCs and (D) hfVM following IFNγ stimulation. (E-I) Comparison of cell types post treatment with IFNγ. (E) Clustered heatmap displaying immune-related pathways expressed by cells following IFNγ stimulation. (F) Differential expression plot comparing IFNγ treated-NPC to IFNγ treated-hfVM. (G-H) Genes expressed at a significantly higher (G) or lower (H) level by IIFNγ treated-NPCs than IFNγ treated-hfVM. (F) Differential expression plot comparing IFNγ treated-NPC to IFNγ treated-hESC/DAn. Data are represented as mean ± STDEV. One way ANOVA; n.s. = not significant *p<0.05, **p<0.01, ***p<0.005, ****p<0.001.

In keeping with the surface protein expression, nCounter analysis confirmed expression of *HLA-A*, *-B*, and *-C*, with ESCs and DAn expressing higher levels of *HLA-A* and *HLA-C* than the NPCs and fVM, and low expression of the MHC-II encoding genes. *HLA-DPB1* was the most constitutively expressed Class-II gene across all cell types, alongside very low levels of *HLA-DPA1* and *-DMA*. Constitutive *CD40* expression was only above the limit of detection for hESC. No cell types expressed *CD80* or *CD86.* Expression levels of these key genes was comparable for NPCs and hfVM tissue (Figure 2A).

Gene (Sup Figure 3F-G) and pathway (Figure 2B) clustering analyses, performed based on the entire gene set, revealed that the hfVM, and NPCs and DAn in particular, are transcriptionally similar, while the hESCs appeared distinct. Differential gene expression analysis revealed 45 upregulated and 22 down regulated genes (significance = p value <0.01) in NPCs compared to cells dissociated from the hfVM (Figure 2C) as well as changes in immune-relevant gene expression through their differentiation from hESC to NPC to DAn (Figure 2D).

Genes significantly more highly expressed in NPCs compared to hfVM were largely anti-inflammatory in nature, including co-inhibitory molecules and anti-inflammatory mediators (*CD112*, *CD276, CSF1, CTNNB1*), TFGβ pathway genes (*TGFB1, TGFBR2*) and non-canonical MHC-I genes such as *HLA-E* (Figure 2E). Whereas genes more highly expressed by hfVM were more pro-inflammatory, including chemokines (*CXCR4, CCL26*), complement and antiviral response genes (*C5, MX1*) and genes commonly expressed by antigen presentation cells (*CD14, CTSS*) (Figure 2F), together suggesting that at the gene level, hfVM cells appear more pro-inflammatory in nature than the NPCs.

Performing differential gene expression analysis across the different cell differentiation stages revealed greater expression of TFGβ pathway associated genes in NPCs compared to hESC (*TGFB1, TGFBR1, TGFBR2*), with expression of *TGFBR1* also being significantly higher in NPCs than in DAn. NPCs and DAn also expressed higher levels of co-inhibitory molecules *CSF1, CD276, CD112* and *CD113* than the hESC, with *CSF1* and *CD112* expression being greatest at the day 16 NPC stage (Figure 2G). Furthermore, several pro-inflammatory genes were found to be expressed at lower levels by the NPCs compared to the hESC and/or DAn, including genes involved in antigen processing (*PSMB8, PSMB9, TAPBP*) which were lower in the NPCs than hESC and DAn, except *TAPBP* in which expression levels correlated with differentiation stage. Complement genes showed opposing directionality, with *C1QBP* decreasing and *C4A/B* increasing with differentiation, and other pro-inflammatory genes were expressed at a lower level by NPCs than the hESC or DAn (*CCL26, MX1*) (Figure 2H). Several genes differentially expressed between hESC, DAn and the day 16 NPCs were related to cell cycle, division and neuronal cell function, as would be expected as the cells transition from a replicative to a terminally differentiated state.

Together these data show that at baseline, the NPCs appear to express higher levels of genes encoding molecules with anti-inflammatory properties and lower levels of pro-inflammatory genes, when compared to hfVM tissue, and that the NPC stage of differentiation also appears to be less immunogenic than the hESC or mature DAn, though the differences between NPC and DAn states were minimal.

### hfVM and NPCs both increased expression of antigen processing and presentation genes in response to IFN ***γ*** exposure

In response to IFNγ exposure, the NPCs upregulated 43 genes (Figure 3A). Of these, 31 were also upregulated in hfVM (Figure 3B) and 25 in DAn (Sup Figure 3C) (of a total of 37 and 35 genes, respectively), showing similar responses to the inflammatory stimulus. As expected, many of these were IFN-response genes, including *IRF1* and *GBP-1* and *-5*, showing that NPCs, DAn and hfVM tissue have the capacity to respond to IFNγ. The majority of genes significantly upregulated by NPCs, hfVM and DAn were those involved in antigen processing and presentation pathways, many of which were below the limit of detection in the “no treatment” (NT) condition (Figure 3C-D, Sup Figure 3D), including (*TAP-1-2, TAPBP, PSMB8-10*) and MHC-I antigen presentation genes (*HLA-A, -B, -C, B2M*). DAn did not upregulate *HLA-A, -B*, or -*C* in response to IFNγ, instead only upregulating non-canonical MHC-I gene *HLA-E* (Figure 3A-B; Sup Figure 3C). However, baseline differential expression analysis showed that DAn express significantly higher levels of these genes without exposure to inflammatory signals (Figure 2A). NPCs and hfVM also increased expression of *TLR3* and *HVEM* following IFNγ exposure, two common pro-inflammatory receptors involved in the antiviral response.

NPCs, hfVM, and DAn also all upregulated several anti-inflammatory genes in response to IFNγ, including *CD274* (encoding co-inhibitory molecule PD-L2) and *IDO-1* (responsible for catabolism of tryptophan to anti-inflammatory kynurenine). Only 6 genes were uniquely upregulated by hfVM and not NPCs, 2 of which were T-cell chemoattractants *CXCL9* and *CXCL10* (Figure 3B). Notably, the two genes with the highest upregulated fold-change by DAn were *CXCL10* and *CXCL11* (Sup Figure 3C-D).

In line with previous reports^53^, hESC did not show any transcriptional response to IFNγ (Sup Figure 3E), despite expression of the IFN receptor *IFNGR1,* resulting in further separation of the ESC cluster on pathway cluster analyses of IFN-treated cells (Figure 3E).

Following IFNγ exposure, the number of genes significantly differentially expressed between NPCs and hfVM increased from 69 (without treatment) to 82 (Figure 3F). 65% of the genes were retained from the untreated comparison, while additional genes, more highly expressed by IFNγ treated-NPCs than IFNγ treated-hfVM, included pro-inflammatory genes associated with TNF pathway activation (*TRAF3, TRAF5, TNFSF11*) and caspase activation (*CASP1, CASP8*), perhaps suggesting that NPCs may be more susceptible to inflammation induced cell death than hfVM tissue (Figure 3G). IFNγ treated-hfVM was found to express significantly higher levels of complement genes (*C3*), antigen presentation markers (*CD83, HLA-DRA, HLA-DPA1*) and other proinflammatory genes, such as *CCL3* (Figure 3H) when compared to IFNγ-treated NPCs. While the differentially expressed genes between IFNγ treated-NPC and IFNγ treated-DAn remained similar to those without IFNγ treatment. The IFNγ treated-hESC were of course strongly separated from both of their lineage committed progeny (Figure 3I).

### NPCs do not induce T-cell activation or proliferation in vitro but are immunosuppressive

Next, all cell types (with and without pre-exposure to IFNγ) were tested in three *in vitro* assays involving their co-culture with (A) magnetic activated cell sorting (MACS)-enriched, proliferation dye-labelled T-cells, (B) proliferation dye-labelled PBMCs and (C) proliferation dye-labelled T-cells and monocyte-derived dendritic cells (moDCs) autologous to the T-cells. Successful generation of moDCs was confirmed via assessment of their morphology, surface expression of key molecules and by demonstrating their ability to engulf and present antigen to

T-cells inducing T-cell activation (Sup Figure 4A-D). Allogeneic and autologous moDCs were also used as positive and negative controls in the T-cell co-culture assays (Sup Figure 4E).

**Figure 4.**
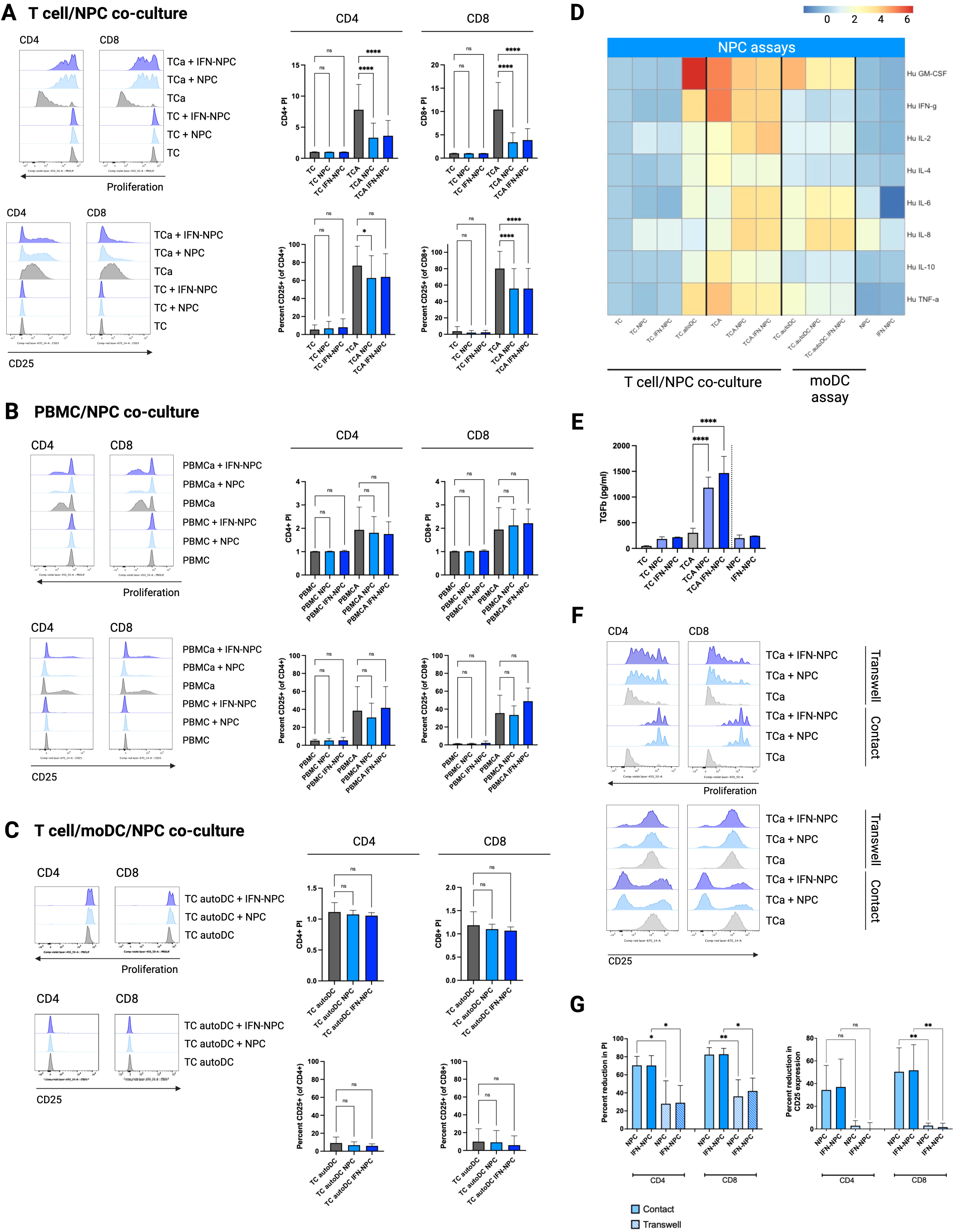
NPCs do not induce T-cell activation or proliferation in in vitro immunogenicity assays, instead they display largely contact-dependent immunosuppressive properties. (A-C) Representative histograms and quantification of T-cell activation represented by proliferation and CD25 expression; untreated and IFNγ treated-NPCs were tested in the T-cell co-culture (A; n=16), PBMC co-culture (B; n=5), and T-cell/moDC co-culture (C; n=10). Quantification of proliferation index (PI) using Modfit modelling software, and CD25 expression in CD4+ and CD8+ T-cells from all assays. Supernatants from the NPC co-cultures with T-cell and T-cell/moDC or NPC/IFNγ treated-NPC only wells were collected and subjected to the 8-plex human cytokine Luminex panel, measuring concentration of IFNγ, TNFα, IL-2, IL-6, IL-8, IL-4, IL-10, GM-CSF (D) or TGFβ ELISA (E). Heatmap displays cytokine concentration relative to untreated T-cell only well. N=3. (F-G) The 5-day NPC-T-cell co-culture assay was repeated as above (contact) or in a transwell format, utilising 0.4 μm transwell inserts separating T-cell and NPC populations, then T-cell proliferation and CD25 expression were measured (F) and quantified (G) N=3. Data are represented as mean ± STDEV. One-way ANOVA; n.s. = not significant *p<0.05, **p<0.01, ***p<0.005, ****p<0.001.

Assay readout was T-cell proliferation and expression of the T-cell activation marker CD25, and in a subgroup of experiments cytokine production, at day 5. No significant T-cell proliferation, activation or cytokine production was seen, in any of the assays, in response to NPCs – even after pre-exposure to IFNγ (Figure 4). Proliferation, activation and production of GM-CSF, IFNγ and TNFα cytokines was observed following T-cell co-culture with allogeneic moDCs – confirming the ability of this assay to detect T-cell allo-responses – and polyclonal anti-CD3/28 bead stimulation (TCA) (Figure 4A-D and Sup Figure 4E). In fact, rather than inducing an immune response, NPCs were found to suppress T-cell proliferation, activation, and the production of some proinflammatory cytokines when added to anti-CD3/28 polyclonally stimulated T-cells (Figure 4A,D and Sup Figure 5A). Furthermore, the concentration of the immunosuppressive cytokine TFGβ was significantly higher in co-cultures of activated T-cells with NPC or IFN-NPCs (Figure 4E). NPCs were also observed to suppress T-cell responses to alloDCs, although somewhat variably across donors (Sup Figure 5D-F).

**Figure 5.**
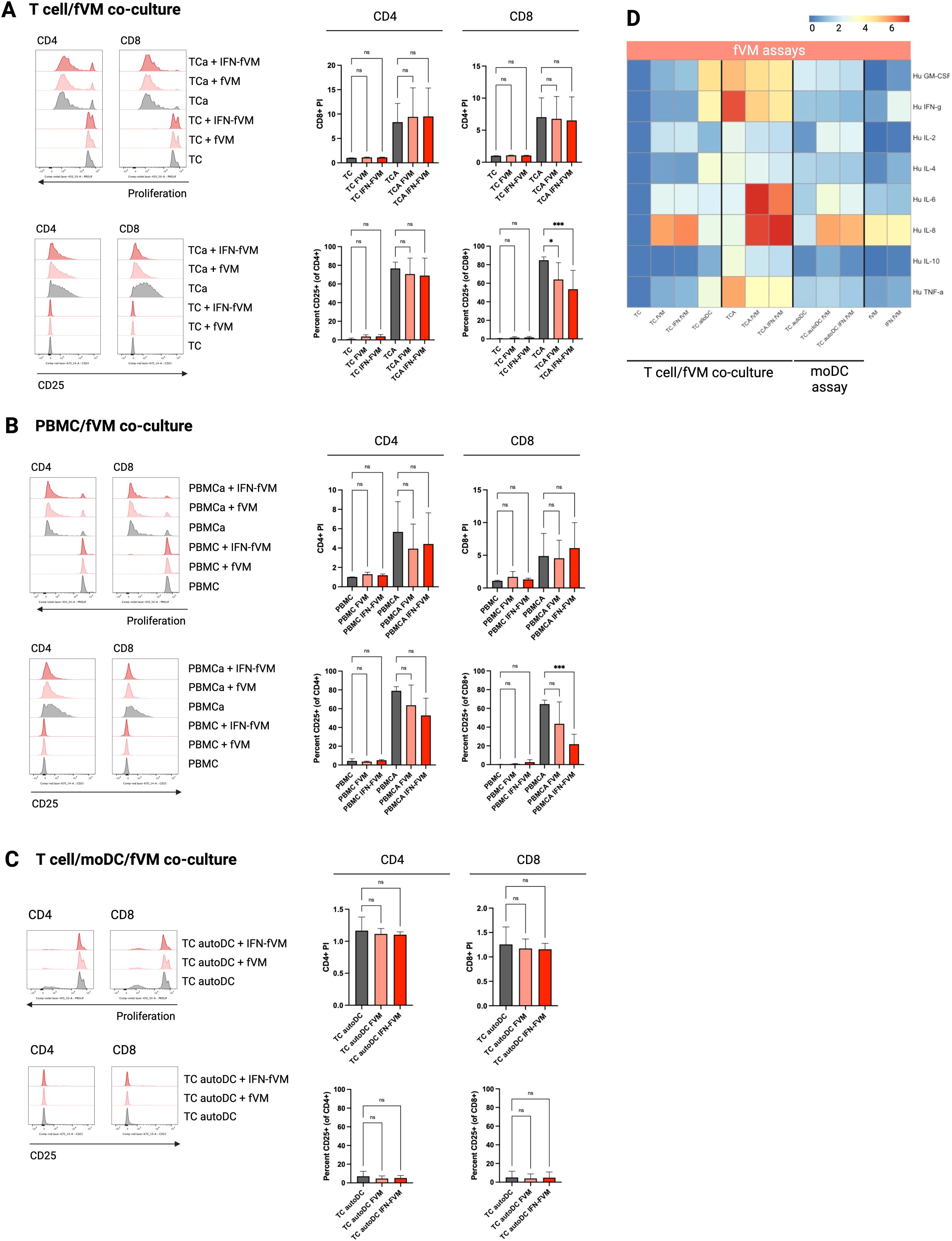
hfVM are neither immunogenic nor immunosuppressive in vitro. (A-C) Representative histograms and quantification of T-cell activation represented by proliferation and CD25 expression; untreated and IFNγ treated-hfVM were tested in the T-cell co-culture (A; n=8), PBMC co-culture (B; n=5) and T-cell/moDC co-culture (C; n=5). Quantification of proliferation index (PI) using Modfit modelling software, and CD25 expression in CD4+ and CD8+ T-cells from all assays. (D) Supernatants from the hfVM co-cultures with T-cell and T-cell/moDC or hfVM/IFNγ treated-hfVM only wells were collected and subjected to the 8-plex human cytokine Luminex panel, measuring concentration of IFNγ, TNFα, IL-2, IL-6, IL-8, IL-4, IL-10, GM-CSF. Heatmap displays cytokine concentration relative to untreated T-cell only well. N=3. Data are represented as mean ± STDEV. One-way ANOVA; n.s. = not significant *p<0.05, **p<0.01, ***p<0.005, ****p<0.001.

### NPC mediated suppression is largely contact dependent

To understand the importance of direct cell contact in NPC-mediated T-cell suppression we repeated the T-cell/NPC co-culture assay using a transwell. This revealed significantly reduced suppression of T-cell activation and proliferation when the cells were physically separated suggesting that suppression is largely contact dependent (Figure 4F-G).

To examine the immunosuppressive mechanism, we screened our nCounter dataset for potentially immunosuppressive molecules expressed by untreated and IFNγ treated-NPCs, as both had been shown to be immunosuppressive. This revealed high levels of co-inhibitory molecules *CD276* (*B7-H3*), *CD112*, *CD113* and *CD155*, while *CD274 (PD-L1), PDCDLG1* (*PD-L2*), or *VTCN1* (*B7-H4*) were not expressed (Sup Figure 6A). However, neither inhibition of CD276 (neutralising antibody; Sup Figure 6B), nor TIGIT - the receptor of CD112/CD113/CD155 (blocking antibody; Sup Figure 6C) - inhibited NPC-mediated T-cell suppression. CD47 (also referred to as IAP) is a known immunosuppressive molecule that is also used as a lineage defining molecule for our NPC product. Blocking of this molecule also failed to reverse NPC-mediated T-cell suppression (Sup Figure 6G), despite high expression at the gene (Sup Figure 6E) and protein (Sup Figure 6F) level. CD47 is known to have immunoregulatory properties and overexpression is now a widely utilised strategy for generating hypoimmunogenic cell lines for therapy, so despite not being responsible for the suppressive effects of the cells, high constitutive expression of this protein may be beneficial for the survival of this product *in vivo*.

**Figure 6.**
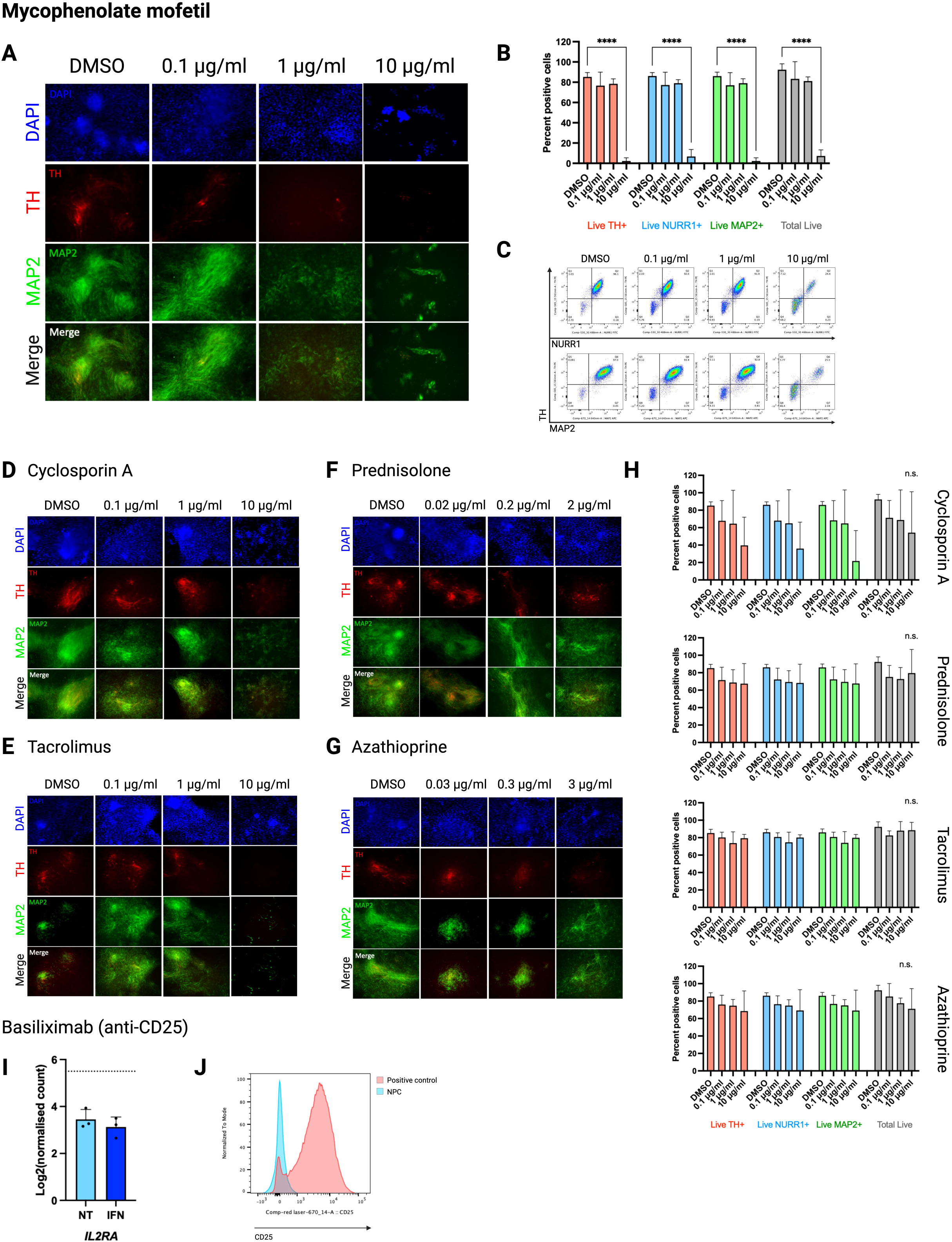
MycophenolatemofetilnegativelyaffectsNPCsurvivalanddifferentiation-implications for clinical trials. Day 16 NPCs were differentiated *in vitro* to day 45+ in the presence of 3 concentrations of (A-C) mycophenolate mofetil, (D) cyclosporin A, (E) tacrolimus, (F) prednisolone or (G) azathioprine. Immunofluorescence (A, D-G) and flow cytometric (B-C, H) analyses were performed to determine the extent of survival and correct lineage differentiation. Representative immunofluorescence images display nuclear stain (blue), TH (red), MAP2 (green) and merged images. Flow cytometry was performed to measure surface expression of NURR1, TH and MAP2 as well as percent live cells using ZombieNIR dye. Flow cytometry analysis was performed to quantify the effect of each immunosuppressive agent on the percent cells positive for each surface marker and the percentage of live cells. (I) To confirm the lack of need to test CD25-targeting Basiliximab, a lack of CD25 expression was confirmed at the gene level (*IL2RA* gene) using nCounter NPC and IFNγ treated-NPC data (I) and at the protein level by flow cytometry compared to a positive control cell type (J). Data are represented as mean ± STDEV. One-way ANOVA; n.s. = not significant *p<0.05, **p<0.01, ***p<0.005, ****p<0.001.

Due to the incomplete inhibition of suppression in the transwell assay, contact-independent mechanisms were also investigated. NPCs did not express genes for anti-inflammatory cytokines *IL-4*, *IL-10*, or *IL-13*, but were found to express high levels of *TGF* β (Sup Figure 6H). We first went on to demonstrate that rhTFGβ is able to suppress polyclonally stimulated T-cells in a dose-dependent manner (Sup Figure 6I) as this has been shown previously as a suppressive mechanism of other hPSC-derived NPCs^54^. However, NPC-mediated suppression of T-cells was not inhibited by the addition of TFGβR1 or TFGβR2 inhibitors (Sup Figure 6J). IDO-1/TDO-mediated catabolism of tryptophan to kynurenine is a well-reported mechanism of suppression in other PSC-derived products^55,56^. While the NPCs did not express *IDO-1* or *TDO* at baseline, they did upregulate *IDO-1* expression following exposure to IFNγ (Sup Figure 6K) so this mechanism was tested, in case IFN-treated NPCs employ a different mechanism of suppression to the untreated cells. Inhibition of IDO-1, using 1-LMT, had no significant effect on suppression however. Finally, as NPCs express (at low level) surface CD39 and CD73 (ectonucleotidases known to convert ATP to immunosuppressive adenosine; Sup Figure 6M) we tested the role of adenosine in NPC-mediated T-cell suppression by blocking the main T-cell adenosine receptor A2A. This did not affect NPC-mediated suppression (Sup Figure 6N). While we were not able to find the immunosuppressive mechanism of the NPCs, our comprehensive search strategy ruled out several known mechanisms.

### hfVM, ESC and DAn are neither immunogenic nor immunosuppressive in vitro

Like NPCs, cells dissociated from hfVM tissue, ESC and DAn did not induce T-cell proliferation or CD25 expression *in vitro* (Figure 5, Sup Figure 7). However, in contrast to NPCs, these cells also did not suppress polyclonally stimulated T-cell proliferation or activation (Figure 5A, Sup Figure 7A,C) suggesting that T-cell suppression is a unique feature of the progenitor cell state.

Co-culture of fVM, IFNγ treated-fVM, DAn and IFNγ treated-DAn led to a slight increase in some supernatant cytokines, including IL-6 and IL-8 production (Figure 5D, Sup Figure 7E) when compared to the NPCs (Sup Figure 5G) again highlighting the low immunogenicity of the NPCs in culture.

### Mycophenolate mofetil negatively affects NPC survival and differentiation - implications for clinical trials

While the *in vitro* data presented here suggest that NPCs are at low risk of immune rejection, immunosuppressive drugs are standardly given in regenerative medicine trials including STEM-PD. Prior to the initiation of the study, the immunosuppressive agents being considered included: steroids (methylprednisolone and prednisolone), basiliximab, tacrolimus (or cyclosporine if not tolerated), mycophenolate mofetil (MMF) and azathioprine. Therefore, prior to the start of the study we demonstrated that NPCs do not express CD25 (the target of basiliximab; Figure 6I-J) and tested the effect of the other immunosuppressive drugs on the survival and differentiation of NPCs to mature DAn *in vitro*. Of the drugs tested, only MMF negatively impacted NPC survival and expression of mature dopamine cell markers TH, NURR1 or MAP2 at the highest concentration tested (selected based on previous *in vitro* studies^57–63^). As a result of this finding, MMF was removed from the STEM PD protocol (NCT05635409), while cyclosporin A, tacrolimus, prednisolone, azathioprine and basiliximab were retained.

## Discussion

Here we have shown that RC17-derived NPCs, generated and handled as per the STEM-PD clinical trial protocol, are not immunogenic *in vitro* but instead possess immunoregulatory properties.

Throughout this project, NPCs were compared to hfVM tissue, selected as fVM has previously been shown to survive long term in the brains of people with Parkinson’s disease, beyond the initial post-transplant period of immunosuppression^1,11,19,29^. Our data indicate that NPCs are at no higher risk of immune rejection than hfVM tissue, and in fact may be at less risk due to their immunosuppressive qualities. We are further reassured by this finding given that all fVM analyses presented here were performed using one donor, whereas in the TRANSEURO and other fVM trials, cells from up to 6 foetuses were utilised per transplant, meaning that an additional degree of alloreactivity would be expected both within the graft and between donor and recipient. Comparison of the immune profile of the cells through their differentiation, from hESC to mature DAn also shows the day 16 NPCs (the stage at which they are transplanted) to be particularly immunologically “inert”, with low expression of immunogenic molecules and high expression of immunoregulatory molecules compared to the hESC and DAn. This is important as the risk of immune rejection is likely to be highest early post-transplant, in the setting of post-surgery induced inflammation and a disrupted blood brain barrier.

To mimic the post-surgical environment and known inflamed environment of the PD brain, we profiled our cells pre- and post-exposure to inflammatory cytokines. Of the cytokines tested, IFNγ induced the most marked effects, even at relatively low concentrations (concentrations in the brain at the site of surgery are unknown). However, while hfVM, NPCs and DAn all upregulated MHC-I antigen processing and presentation genes – they all remained non-immunogenic in our assays, providing reassurance that even in an inflammatory environment, the NPCs are unlikely to induce a significant T-cell response. Although modelling immune responses *in vitro* is known to be very challenging, we were reassured that our assays were capable of detecting T-cell responses (proliferation, activation and cytokine production) in response to other allogeneic cells (allo-moDCs).

Despite their favourable immune profile *in vitro*, due to the highly invasive nature of the surgery and high stakes involved, patients will continue to be immunosuppressed to further reduce the risk of rejection. Meta-analysis of the hfVM trials concluded that graft survival was improved in trials that included immunosuppression for at least one year post engraftment^50^. However, our data suggest that aggressive immunosuppression (based largely on regimens used to prevent solid organ transplant rejection), that are currently being used in patients with PD receiving neural transplantation, may not be required. And that a lower intensity regimen, or one of shorter duration, may be sufficient. This would reduce the well-recognised risks of immunosuppression, particularly risk of infection, in our older patient population. Our work also speaks to the importance of drug selection as here we demonstrate that MMF negatively impacts the survival and differentiation of NPCs *invitro.* This is rarely considered and often neglected in the preclinical work up of regenerative cellular therapies. In response to our data, MMF was switched out for azathioprine in the STEM-PD protocol (NCT05635409).

Aside from immunosuppression, several other approaches are being considered with regards to dampening down immune responses to neuronal and other cell therapies, including the development of “hypoimmunogenic” cell lines, lacking MHC-I and/or MHC-II and overexpressing immunomodulatory molecules such as CD47^64,65^. Co-transplantation of regulatory immune cells is also being explored in autologous transplantation to suppress surgery-induced immune activation^66^. Our data suggest that genetic engineering of RC17-derived NPCs is not required. They constitutively express high levels of CD47, lack MHC-II expression and do not appear capable of directly activating T-cells, even after exposure to inflammatory stimuli. Genetically engineering cells comes with risks such as potential off-target effects and NK activation through the missing-self response. Furthermore, in our case of neuronal therapy, MHC-I itself is known to have roles in synaptic pruning and neurodevelopment^67^, raising further questions around its modification.

In summary, our *in vitro* data suggest that the STEM-PD dopamine cell product is of low immunogenicity and that it is unlikely to induce high levels of immune rejection post-engraftment. Further preclinical assessment of the product, using for example humanised mouse models, will be of value and every opportunity should be made to understand the risks of immune rejection, and the risks of immunosuppression, by carefully assessing and collecting samples from patients undergoing transplantation on the STEM-PD study. However, we suggest that generating MHC-I knockouts is not a necessary step for the development of the next generation of hPSC derived dopamine cell therapies for PD, and that patients may benefit from more refined immunosuppressive regimens moving forward.

### Limitations of the Study

The primary limitation of our study is the relative insensitivity of *in vitro* immune assays and lack of better *in vitro* models for assessing rejection responses, especially over time. Investigating the immunogenicity of RC17-NPCs *in vivo*, by using human immune system (HIS) mice, would provide further reassurance., however these models also have their challenges - including variable levels of engraftment, a limited myeloid compartment and their limited lifespan before they develop graft vs. host disease. This is a particular issue for testing the immunogenicity of progenitor cells that require time to mature and differentiate *in vivo*.

To date, we have focused our analysis on the peripheral immune response to the product, however, as a neuronal therapy it is important also to consider the response of brain resident immune cells - the microglia. Further work to understand the interaction of RC17-NPCs with microglia will be important as previous studies have revealed that microglial activation can be detrimental to stem cell derived NPC product survival and differentiation *in vivo* ^68^ and have shown microglial infiltration in similar allografts post-mortem^69^.

## Acknowledgements

This work was supported by UK-Regenerative Medicine Platform (UKRMP; MR/S020934/1), Horizon 2020-funded Neural Stem Cell-Reconstruct (NSC-R; G101558), Rosetrees Trust and John Black Charitable Fund (G118482) and NIHR Biomedical Research Centre funding to J.L.J. (BRC-1215-20014). A.J.C.’s studentship was supported by Cambridge Trust and Masonic Charitable Trust Foundation. We performed flow cytometric analysis in the Cambridge NIHR BRC Cell Phenotyping Hub (Department of Medicine, University of Cambridge). The views expressed here are those of the author(s) and not necessarily those of the NIHR.

## Author contributions

The authors confirm contribution to the paper as follows: A.J.C., S.F., S.K.H, S.Q., and X.H. performed experiments. A.J.C. and S.Q. analysed data. A.J.C. wrote the draft manuscript, with input from J.L.J and R.A.B. All authors contributed to the design of experiments and data interpretation and approved the final version of the manuscript.

## Declaration of interests

R.A.B has provided consultancy advice around dopamine cell-based therapies to Aspen Neuroscience, BlueRock Therapeutics, FCDI and Novo Nordisk. J.L.J reports consultancy work for Enhanc3DGenomics, Sanofi and Roche. S.Q. is a current employee at Insmed Inc.

## Methods

### Resource availability

Further information and requests for resources and reagents should be directed to and will be fulfilled by the lead contact, Joanne Jones.

### Cell culture conditions

All long-term cultures were performed in the presence of 100 U/ml penicillin-streptomycin (Thermo Fisher Scientific). Standard culture conditions involved the use of Corning tissue culture plates (Thermo Fisher Scientific) and incubation at 37°C/5% CO2.

#### Human Embryonic Stem Cells (hESC) and derivatives

RC17 hESC (Roslin Cells, hPSCreg name: RCe021-A) are a clinical grade, healthy female hES cell line with a normal karyotype derived under GMP and xeno-free conditions, for these experiments research-grade RC17 hESCs were utilised. The cells were obtained from the inner cell mass of a 3-day old blastocyst stage embryo, surplus or unsuitable for clinical use. RC17 has normal pluripotency marker expression and can differentiate to all three germ layers *in vitro*. RC17 hESCs were utilised under study protocol REC No: 21/EE/0051.

Undifferentiated ESCs were routinely cultured in StemMACS iPS-Brew, XF medium (Miltenyi Biotec) on culture plates pre-coated with Laminin-521 (Biolamina) at 0.5 ug/cm^2^. All single cell dissociations were done by incubating with StemPro Accutase (Gibco) for 4-6 minutes at 37°C. Following dissociation, cells were re-plated with 10 uM StemMACS^TM^ Y-27632 ROCK inhibitor (ROCKi; Miltenyi Biotec). Media was changed every 24 hours.

Upon differentiation of hESCs to day 16 mesencephalic dopaminergic (mesDA) neural progenitor cells (NPC), NPCs were cultured in 2% B-27-supplemented Neurobasal CTS Media (plus 2 uM L-glutamine and 0.2% vol/vol penicillin–streptomycin; Thermo Fisher Scientific) plus 20 ng/ml BDNF, 0.2 uM ascorbic acid (AA) and 100 ng/ml FGF8b on culture plates pre-coated with Laminin-111 (Biolamina) at 1 ug/cm^2^. All single cell dissociations were done by incubating with StemPro Accutase (Gibco) for 4-6 minutes at 37°C. Following accutase dissociation, cells were re-plated with 10 uM ROCKi. Media was changed every 48 hours.

Upon differentiation of day 16 NPCs to mature DAn, mature DAn were cultured in the same media as described above for the NPCs, with the addition of 1 uM DAPT (N-[(3,5-difluorophenyl)acetyl]-L-alanyl-2-phenyl]glycine-1,1-dimethylethyl ester; R&D Systems) and 500 uM cyclic AMP (cAMP; Sigma Aldrich), on culture plates pre-coated with Geltrex (Thermo Fisher Scientific) diluted in DMEM (Thermo Fisher Scientific) at 0.01% for one hour at 37°C. Half the media was changed every 48 hours.

#### Human primary foetal ventral mesencephalon (hfVM) collection and maintenance

Fresh human foetal ventral midbrain tissues were dissected from foetuses aged 7-10 weeks post conception collected at pregnancy termination from the Rosie Hospital, Cambridge (ethics ref: 96/085). hfVM tissues were stored in Hibernate-E Medium (Thermo Fisher Scientific) for up to 4 days before washing in DMEM and incubation with Accutase for 30 minutes at 37°C and agitating at 10-15-minute intervals. Accutase was removed and the cells were then resuspended in DMEM with 2% B-27-supplemented 1% FBS and 1% PSF (Thermo Fisher Scientific) and plated on PDL/Laminin-2020-coated culture plates.

### Differentiation of hESC to NPC/DAn

hESC to NPC: RC17-hESC were differentiated to ventral mesencephalic dopaminergic neural progenitors *in vitro* using the 16-day GMP-compliant protocol derived by Nolbrant *et al.* ^32^ Briefly, on day 0, RC17 hESC, from a 70-90% confluent well, were detached with EDTA and plated on Laminin-111 coated plates at a density of 10,000 cells/cm^2^ with 10 uM ROCKi. Cells were cultured in N-2 media supplemented with 10 uM SB431542, 100 ng/ml Noggin, 300 ng/ml Shh-C24II and 0.85 uM CHIR9902 until day 9. Media was changed to N-2 media plus 100 ng/ml FGF8b until day 11, when they were Accutase-dissociated and replated (at 0.8×10^6^ cells/cm^2^) onto Laminin-111 coated plates with 10 μM ROCKi. Cells were then cultured from day 11 to day 16 in 2% B-27-supplemented Neurobasal CTS Media plus 20 ng/ml BDNF, 0.2 uM ascorbic acid (AA) and 100 ng/ml FGF8b. At day 16, mesDA NPCs had formed (quality testing to verify correct identity was performed using quantitative PCR, flow cytometry, and immunofluorescence). Cells were either cryopreserved for use in subsequent experiments or terminally differentiated to mature dopaminergic neurons *in vitro* (see below).

NPC to DAn: From day 16-45, terminal differentiation and maturation of these cells was performed also as per Nolbrant *et al.* ^32^; cells were replated at a density of 250–400,000 cells/cm^2^ on plates pre-coated with 0.01% Geltrex in DMEM for one hour at 37°C. They were cultured until at least day 45 in B-27-supplemented Neurobasal CTS Media with 20 ng/ml BDNF, 0.2 uM ascorbic acid (AA), 100 ng/ml FGF8b, 1 uM DAPT and 500 uM cAMP.

### Treatment of cells with inflammatory stimuli

hESC, NPC, DAn or hfVM were seeded at a density of 0.5×10^6^ cells/cm^2^ on 24 well plates (for Nanostring experiments) or 96 well plates (for *in vitro* assays) with their respective media and coatings described above. For initial optimisation experiments, (Sup Figure 1) cells were treated for 0-72 hrs with 0-50 ng/ml IFNγ, TNFα, IL-1β or combinations of two or all three at 37°C. Once the duration and concentration of inflammatory stimuli was selected, all further treatments were done for 24 hrs with 5 ng/ml IFNγ and/or TNFα.

### Immunofluorescence staining of ESC, NPC and DAn

All steps were performed at room temperature unless otherwise stated. NPCs (d16) or mature DAn (d45+) were first fixed with a final concentration of 4% paraformaldehyde (PFA; Sigma Aldrich). For DAn, media could not be completely aspirated to avoid lifting cells from the plate. Instead, for fixation and all washes, several half volume washes were performed to dilute the media. Cells were then washed with PBS then blocked in blocking buffer (0.25% Triton + 5% donkey serum) for 1 hr. Cells were washed with PBS-Tween then stained with primary antibodies (see key resource table) diluted in blocking buffer for 1 hour at room temperature then overnight at 4°C. Cells were washed three times with PBS then incubated with secondary antibodies for 1 hour (see key resource table). Cells were washed three times further, then stained with DAPI (41Z,6-diamidino-2-phenylindole; 1:1000; EMD Millipore) in blocking buffer for 10 minutes before three final washes. Images were acquired using an EVOS FL fluorescent imaging microscope (Life Technologies or Leica).

### Flow cytometry of hESC, NPC, DAn and hfVM

ESCs and NPCs were thawed and recovered in complete RPMI media then seeded and washed with PBS. DAn and fVM were washed with PBS. All cells were then left untreated or treated with inflammatory stimuli for 24 hrs. After 24 hrs, media was removed, cells collected using Accutase then washed with PBS. Cells were stained for viability using Zombie NIR or Zombie Aqua Fixable Viability kits (BioLegend) for 15 minutes in the dark at 4°C. Live/dead staining was stopped with FACS buffer (2.5% FCS in PBS) and centrifuged at 400*g* for 6 minutes. Cells were then blocked on ice with PBS + 7.5% FCS for 15 minutes before staining with antibody cocktails (see key resource table) containing TruStain Monocyte Blocker (Biolegend) in FACS buffer for 45 minutes at 4°C. Fluorescence single stains for compensation were performed using single stained cell samples and/or UltraComp eBeads^TM^ (Invitrogen). Cells were washed then run fresh or fixed in 0.25% PFA in PBS. Flow cytometry was performed on an LSR Fortessa or LSR-II Analyser (BD Biosciences). All flow cytometry data was analysed using FlowJo software (BD Biosciences).

### RNA extractions for qPCR and nCounter

Cells with or without treatments (5 ng/ml IFNγ, TNFα or both), were resuspended in RLT buffer and stored at -80°C until use. RNA was extracted using the RNeasy® Plus Mini Kit (Qiagen) according to the manufacturer’s instructions. RNA quality and quantity was confirmed using the Agilent 2100 Bioanalyzer then frozen at -80°C until used for qRT-PCR or Nanostring nCounter.

### Quantitative reverse transcriptase-polymerase chain reaction (qRT-PCR)

0.5-1 ug of RNA was reverse transcribed to cDNA using the High-Capacity cDNA Reverse Transcription Kit (Invitrogen). Following this, a SYBR-green qPCR reaction was performed with gene-specific primers (Sigma) as per Nolbrant *et al.* ^32^ in triplicates in a 384-well plate (MicroAmp; Life Technologies) followed by amplification by qPCR in the Quantstudio 12K Flex qPCR machine (Life Technologies). *GAPDH* and *ACTB* were used as the reference genes for all samples. qPCR data was analysed using average cycle threshold (CT)-values in the Thermo Fisher ConnectTM cloud software (Thermo Fisher) and all data are represented using the double-delta CT method.

### Nanostring nCounter

Three distinct vials of RC17 hESCs, differentiated in separate experiments to day 16 NPC and day 45 DAn, and hfVM from three donors, were cultured for 24 hrs without treatment or with IFNγ, TNFα or both (5 ng/ml) and RNA extracted and the quality and quantity of this was confirmed. Nanostring nCounter was performed according to the manufacturer’s instructions using the Human Immunology V2 Panel with a custom Panel Plus of 20 genes (see key resource table). Briefly, 100 ng of extracted RNA was hybridised for 20 hours at 65°C with two 50 nucleotide probes: a biotinylated capture probe and an oligo-tagged reporter probe. After binding to the RNA, the hybridised mixture was loaded into a streptavidin-coated cartridge causing the biotinylated-capture probe to bind to the cartridge surface. After washing to remove excess probes, an electrical current was applied to immobilise and align the RNA-probe hybrids for digital imaging (MAX/FLEX system). 555 fields of view (FOV) were imaged per sample. Gene expression was normalised to the geometric mean of housekeeping genes using the nSolver software and a threshold for limit of detection was determined by 3*mean of negative controls. Data quality and normalisation were checked using NACHO (NAnostring quality Control dasHbOard). Downstream analyses were performed using R and visualised using R packages EnhancedVolcano and Heatmap2.

### Immunogenicity assays

#### Blood collection, PBMC isolation and cell type enrichment/sorting

Blood was drawn by clinicians from healthy volunteers with informed consent in compliance with NHS Health Research Authority ethical approval reference no. 21/EE/0051. All donors were aged between 21 and 63. Exclusion criteria excluded individuals donating blood within 1 month of vaccination or any individual with diagnosed immune-related or autoimmune disorders. Peripheral blood mononuclear cells (PBMCs) were freshly isolated from heparinized blood using Ficoll-Paque Plus (GE Healthcare) density gradient centrifugation.

#### Magnetic activated cell sorting (MACS) enrichment

Desired cell types (Pan T-cells and monocytes for moDC differentiation) were enriched from total PBMCs using magnetic activated cell sorting (MACS). CD14+ monocytes were first positively selected using CD14+ MicroBeads, then T-cells were negatively selected from the CD14-subset using the PanT-cell isolation kit (Miltenyi Biotech) as per the manufacturer’s instructions.

Briefly, whole PBMCs were incubated with CD14+ microbeads for 15 minutes at 4°C then washed and resuspended in 500ul of MACS buffer (0.5% BSA and 2 uM EDTA in PBS). Resuspended cells were then passed through LS columns in the MidiMACS separator (Miltenyi Biotech) and the columns were washed with 9 ml of MACS buffer. CD14-cells were collected for PanT isolation (see below), then the LS columns were removed from the magnetic separator and CD14+ cells were flushed out of the magnetic column, collecting monocytes. Finally, CD14+ cells were washed before being plated for differentiation (see moDC differentiation below).

CD14-cells were washed then resuspended in MACS buffer and incubated for 5 minutes at 4°C with the PanT-cell biotin antibody cocktail (cocktail of biotin-conjugated monoclonal antibodies against CD14, CD15, CD16, CD19, CD34, CD36, CD56, CD123, and CD235a). Additional MACS buffer was added, then cells were incubated for 10 minutes at 4°C with the PanT-cell MicroBead cocktail (magnetic microbeads conjugated to monoclonal anti-biotin antibody (isotype: mouse IgG1) and monoclonal anti-CD61 antibody (isotype: mouse IgG1)). Cells were then passed through LS magnetic columns held on the MidiMACS separator. The column was washed with a further 3 ml MACS buffer and unlabelled T-cells were collected. The purity of the enriched populations was measured using flow cytometry.

#### T-cell staining with eFluor cell proliferation dye

MACS-enriched Pan T-cells and PBMCs were stained with the eBioscience eFluor 450 Cell Proliferation Dye (Invitrogen) according to the manufacturer’s instructions. Briefly, cells were resuspended in PBS and then the cell proliferation dye (diluted in PBS) was added dropwise while agitating the cells to a final concentration of 5 uM, then incubated at 37°C for 10 minutes. Following incubation, cells were washed with 4X volume of ice-cold RPMI + 10% FCS and incubated on ice for 5 minutes. Cells were then washed three times in RPMI + 10% FCS, aliquoted and cryopreserved.

#### Monocyte-derived dendritic cell (moDC) differentiation

All monocytes were utilised fresh due to the known issues of poor viability of cryopreserved monocytes. Full detailed protocol can be found on protocols.io^66^. Monocytes, MACS-enriched from fresh PBMC were seeded at 0.5×10^6^ cells/cm^2^ in RPMI plus 5% heat-inactivated human AB serum (Merck Life Sciences) and 100 U/ml penicillin-streptomycin and cultured with 50 ng/ml rhIL-4 (Biolegend) and 50 ng/ml rhGM-CSF (Biolegend) for 6 days with half media changes on day 3. On day 6, the cells were activated with 100 ng/ml lipopolysaccharide (LPS; Thermo Fisher Scientific) for 24 hours, then washed twice with PBS to remove residual LPS before co-culture.

Successful differentiation to a moDC phenotype was confirmed by morphology and by flow cytometry to determine the presence of mature DC markers (MHC-I, MHC-II, CD80, CD86) and the absence of the monocyte marker CD14. Phagocytic function was demonstrated using 1 um yellow-green FluoSpheres (Thermo Fisher Scientific) and antigen presentation function was demonstrated by successful activation of T-cells following 24-hour moDC pre-exposure to the hyper-immunogenic hemocyanin from Megathura crenulata (keyhole limpet hemocyanin; KLH; Sigma Aldrich).

#### In vitro co-culture assays

Cryopreserved, eFluor 450-stained PBMCs or T-cells were thawed and recovered in RPMI with 5% human AB serum and 100 U/ml penicillin-streptomycin (complete RPMI medium). Cells were plated at a density of 0.5×10^6^ cells/cm^2^ (150,000 cells/well) and co-cultured with the cell of interest (ESC, NPC, DAn, fVM), also at a density of 0.5×10^6^ cells/cm^2^ in a 1:1 media mix of complete RPMI medium and respective cell media (see section 2.1; RPMI-mixed media). Cells were co-cultured for 5 days with half media changes every 48 hours unless stated otherwise. At the end of the 5-day co-cultures, supernatants were collected for ELISA/Luminex analysis and PBMCs or T-cells were collected and stained, and T-cell proliferation and activation measured using flow cytometry (see below). Flow cytometry was performed on an LSR Fortessa or LSR-II Analyser (BD Biosciences). All flow cytometry data was analysed using FlowJo software (BD Biosciences) and quantification of proliferation was performed using Modfit modelling software. All quantified data are presented as mean ± standard deviation (STDEV). Biological replicates (n) are stated in figure legends.

#### PBMC and T-cell co-culture assays

hESC, NPCs, DAn or hfVM were plated 24 hours prior to co-culture on coated 96 well plates at a density of 0.5×10^6^ cells/cm^2^ (150,000 cells/well) and cultured for 24 hours with or without IFNγ (5 ng/ml; see section 2.4). On the first day of co-culture, cells were washed with PBS, and eFluor 450-stained PBMCs or T-cells were seeded. Cells were co-cultured in RPMI-mixed media for 5 days, with half media changes every 48 hours. Several positive controls were utilised: (1) polyclonal T-cell simulation with soluble αCD3 (0.5 μg/ml; clone OKT3; Biolegend) and α-CD28 (1μg/ml; Invitrogen) or Dynabeads™ Human T-Activator CD3/CD28 for T Cell Expansion and Activation (1:30 bead:T-cell ratio; Gibco) or (2) allogeneic moDCs.

#### Monocyte-derived dendritic cell (moDC)/T-cell co-culture assay

MoDCs were differentiated from monocytes and activated as described above. hESC, NPCs, DAn or hfVM were plated 24 hours prior to co-culture on coated 96 well plates at a density of 0.5×10^6^ cells/cm^2^ (150,000 cells/well) and cultured for 24 hours with or without IFNγ (5 ng/ml). On the day of co-culture, cells were washed with PBS, collected using EDTA and seeded into the moDC wells in the presence of 10 μM ROCKi at a density of 0.5×10^6^ cells/cm^2^ (150,000 cells/well). Simultaneously, eFlour 450-stained T-cells were thawed, recovered in complete RPMI media, and seeded at 0.5×10^6^ cells/cm^2^ (150,000 cells/well). The three cell-types were cultured in RPMI-mixed media for 5 days with half media changes every 48 hours. The KLH antigen presentation acted as a positive control for this assay.

#### Transwell assay

The transwell assay was performed in a 24-well plate format. Minimum cell number for effective activation was found to be 500,000 cells/well. NPCs were plated 24 hours prior to co-culture on Laminin-111 coated 24 well plates at 500,000 cells/well and cultured for 24 hours with or without IFNγ (5 ng/ml; see section 2.4). On the first day of co-culture, cells were washed with PBS, following which eFluor 450-stained PBMCs or T-cells were seeded either directly into the well (contact well) or into a 0.4 um transwell membrane insert (Corning Falcon). In contact wells, 1 ml total media was added (1:1 NPC/T-cell media). In transwell wells, 500 ul media was added to the plate, and 500 ul to the transwell insert, thus ensuring that each 1:1 NPC/T-cell had a total volume of 1 ul of media. The assay was performed as previously described with half media replenishment from the plate and transwell insert each 48 hours.

#### Inhibitors and blocking antibody treatments

To test for mechanisms of NPC-mediated suppression, T-cell co-culture assays were performed in the presence of inhibitors and blocking antibodies. In all cases, target cells (NPCs and/or T- cells) were exposed to inhibitors or blocking antibodies for 2 hours prior to the start of the assay then refreshed throughout the duration of the 5-day co-culture. The following functional grade blocking/neutralising antibodies were used, all at 10 μg/ml: monoclonal anti-CD276 (clone MIH35; eBioscience), Ultra-LEAF™purified monoclonal anti-TIGIT Antibody (clone A15153A; Biolegend), monoclonal anti-CD47 (clone B6H12; eBioscience). TGFβR1 inhibition was performed using the SB 431542 inhibitor (Biotechne) at 10 μg/ml and TGFβR1/2 blocking was done using LY2109761 inhibitor (Stratech) at 10 μg/ml. Cells were treated with 200 μM of 1-methyl-L-tryptophan (1-LMT; IDO-1 inhibitor), 1-methyl-D-tryptophan (1-DMT; 1-LMT enantiomer, both Merck Life Sciences) for exploration of a possible tryptophan mechanism of immunosuppression. For adenosine pathway investigation, cells were treated with 2.5 μM concentration of ZM241385 (Adenosine 2A receptor inhibitor).

### Enzyme-linked immunosorbent assay (ELISA)

The Human TGF-beta 1 Duoset ELISA was utilised with the Ancillary reagent kit 1 (Biotechne) to measure secreted TGFβ1 in the supernatants of the T-cell and moDC assays. Briefly, 150 ul supernatants were collected at the end of the 5-day assay, centrifuged at 1500*g* for 10 minutes, then the liquid collected and frozen at -80°C until use. Supernatants were thawed on ice then pre-activated using the sample activation kit of 1.2N NaOH for activation and 0.5M HEPES for neutralisation (specific to the TGFβ kit to activate latent TGFβ). Sample titrations were performed to determine the optimal dilution ensuring absorbance fell within the linear region of the standard curve.

All steps were performed at room temperature unless otherwise stated. ELISA plates were coated with capture antibody (2 ug/ml) overnight, then washed three times with wash buffer (0.05% Tween-20 in PBS). Plates were then blocked for 1 hour using the reagent diluent (1% BSA in PBS) before being washed three times. 100 ul of diluted sample, or standard was added to the coated plate and incubated for 2 hours before washing. Detection antibody (50 ng/ml) was added and incubated for 2 hours before washing again. Colorimetric readout was measured using a streptavidin-horseradish peroxidase (HRP) system: streptavidin-HRP was added to the wells for 20 minutes, washed then substrate solution (1:1 mixture of Color Reagent A (H_2_O_2_) and Color Reagent B (Tetramethylbenzidine)) added for a further 20 minutes. Stop solution (2 N H2SO4) was then added and the optical density of wells measured using a microplate reader at 450 nm (with 570 nm wavelength correction). Final quantification and statistics were performed on the corrected wavelength value (450 nm – 570 nm) and the concentrations of samples interpolated from the standard curve generated using known standards (concentration range: 31.2-2000 pg/ml) and GraphPad Prism.

### Luminex

The human cytokine Luminex 8-plex assay (Biorad) was utilised to measure the concentration of IFNγ, TNFα, IL-2, IL-4, IL-6, IL-8, IL-10 and GM-CSF in supernatants from T-cell and moDC assays. Supernatants were thawed on ice. All steps were performed at room temperature and all wash steps were performed using a magnetic plate stand unless otherwise stated. Briefly, 50 ul of magnetic beads were added to the Luminex plate then washed twice. 50 ul diluted samples or standards were added to each well and incubated on a shaker for 30 minutes before washing three times. Biotinylated detection antibody was added and incubated on a shaker for 30 minutes before washing three times. 50 ul streptavidin-PE was added and incubated on a shaker for 10 minutes before washing three times. Finally, the sample was resuspended in an assay buffer and mixed for 30 seconds before data acquisition using the Bio-Plex 200 system (Biorad). Sample concentrations were interpolated from the standard curve generated using the standards. Samples were run in technical duplicates and percent coefficient of variation (CV) calculated based upon the equation CV=(s/m)*100, where s= standard deviation of replicates and m= mean of replicates. Any sample with %CV>25 was excluded. Inter-plate variability was calculated by measuring the CV of one inter-plate control sample; inter-plate CVs for each cytokine were as follows: GM-CSF 1%, IFNγ 4.9%, IL-2 13.5%, IL-4 6.7%, IL-6 10.5%, IL-8 0.3%, IL-10 1.1%, TNFα 10.5%. As all inter-plate CVs fell below 25%, no inter-plate correction was performed.

### In vitro differentiation and maturation of DA neurons with immunosuppressive drugs

DA neurons were terminally differentiated and matured *in vitro* as described above (Nolbrant *et al.*^32^) in maturation media supplemented with immunosuppressive agents – Cyclosporine A (Sigma), Prednisolone (Thermo Fisher), Azathioprine (Sigma), MMF (Sigma), or Tacrolimus (Sigma); three different concentrations for each drug were used (see key resource table). Cells were replated at a density of 250– 300,000 cells/cm^2^ on plates pre-coated with Geltrex and cultured until at least day 45 with 75% media changes twice a week with the media containing immunosuppressive drugs. Control cells were treated with only DMSO (Miltenyi Biotec) which was used as a drug diluent.

The mature neurons, post day 45 in culture, were first harvested using Accutase and counted to determine cell survival using the Countess™ II Automated Cell Counter. Flow cytometry and immunofluorescence analyses were performed as described above.

### Analysis, statistics, and visualisation

All statistical analyses and visualisation were performed using GraphPad Prism or R packages. Statistical tests are stated in all figure legends and R packages are stated in results sections where necessary. All figures and graphics were generated using BioRender.

