## Supplementary material for "The immunological profile of RC17 hESC-derived dopaminergic neural progenitor cells *in vitro*: implications for the STEM-PD clinical trial": SupplementalMaterialforSubmission.html

 


Supplemental Figure 1. Generation of the STEM-PD product (RC17 hESC-derived NPCs). RC17 hESCs were differentiated to NPCs over 16 days (A-D) and then further matured DAn over 45 days in vitro (e-G). The correct lineage was confirmed multimodally with hfVM tissue being used for comparison. (A) Immunofluorescence staining of day 16 NPCs for positive differentiation markers LMX1A (green) and FOXA2 (red). (B) hESCs, NPCs and hfVM were subjected to a qPCR panel of markers for progenitor state, patterning, pluripotency, predictors of lineage and mature DAn. (C-D) hfVM and NPCs were examined using flow cytometry for surface expression of positive (FOXA2, IAP, EN1) and negative (PAX6, OCT3/4) markers of differentiation (C) and percent positive cells quantified (D). (E) Immunofluorescence staining of day 45 DAn for positive differentiation markers TH1 (green), GIRK2 (green), TUJ1 (red) and FOXA2 (red). (F-G) DAn were examined using flow cytometry for positive markers of differentiation (F; NURR1, TH, MAP2) and percent positive cells quantified (G).   Data are represented as mean ± STDEV. One way ANOVA; n.s. = not significant \*p<0.05, \*\*p<0.01, \*\*\*p<0.005, \*\*\*\*p<0.001.

Supplemental Figure 2. MHC-I and MHC-II surface expression in response to time course and dose-testing of inflammatory stimuli.  (A) NPCs were treated for 24 hours with 0, 1, 5, or 25 ng/ml IFNg, TNFa and IL-1b alone or in combination and surface expression of MHC-I and MHC-II measured using flow cytometry to determine the minimum concentration needed for expression without inducing cell death. (B) IFNg and TNFa treatments were then tested with a wider dose range and MHC-I/II expression measured. (C) Duration of treatment was then examined; cells were left untreated (green; NT), treated with 5 ng/ml IFNg (red), TNFa (blue) or both IFNg and TNFa (orange) for 0-24hrs and expression of MHC-I, MHC-II and the costimulatory molecules CD80, CD86 and CD40 was measured. Cells were treated for a longer duration from 16-72hrs and MHC-I and MHC-II expression measured. All representative data are displayed as overlayed histograms.

Supplemental Figure 3. nCounter gene and pathway analyses revealed similarities between NPCs, hfVM and DAn. (A) Pathway annotations from the genes in Human Immunology V2 panel plus custom add-on panel (B) Pathway scoring for all cell types combined to validate protein expression finding that IFNg induces a stronger inflammatory response than TNFa. (C/E) Differential expression plot comparing (C) DAn and (E) hESCs stimulated with 5 ng/ml IFNg for 24 hours against baseline of untreated cell type (D) Violin plot displaying genes significantly upregulated by DAn following IFNg stimulation. (F-G) Hierarchical clustering analysis based on gene expression counts, including all genes above threshold in untreated cells (F) or cells treated with 5ng/ml IFNg for 24hrs (G). Data are represented as mean ± STDEV. One way ANOVA; n.s. = not significant \*p<0.05, \*\*p<0.01, \*\*\*p<0.005, \*\*\*\*p<0.001.

Supplemental Figure 4. moDC phenotypic validation and in vitro immunogenicity assay optimisation (A-D) Confirmation of differentiation of CD14+ monocytes to mature dendritic cells (DC) via (A) surface expression of DC+ markers, CD80, CD86, MHC-I and MHC-II and lack of monocyte marker CD14, (B) morphology of monocytes without differentiation (-IL-4, GM-CSF; left), differentiated but inactive (+IL-4, GM-CSF; middle) and activated DCs (+IL-4, GM-CSF + LPS; right), and assessment of their functional capacities to (C) uptake antigen utilising FluoSphere™ as antigen representative and (D) to present antigen to T cells resulting in their activation. Hyper-immunogenic molecule keyhole limpet antigen was added to T cells only (TC) or T cells seeded with autologous DCs (autoDC), and T cell activation measured. T cells or TC and autoDC + aCD3/CD28 dynabead activation was included as control. (E) Positive controls used for the different in vitro assay conditions. T cells were cultured alone or co-cultured with autologous or allogeneic DCs (autoDC, alloDC, respectively) for 5 days with a 1:1 media mix of T cell media and media of cell of interest. Data are represented as mean ± STDEV. One-way ANOVA; n.s. = not significant \*p<0.05, \*\*p<0.01, \*\*\*p<0.005, \*\*\*\*p<0.001. N=3.

Supplemental Figure 5. Representative dotplots from NPC co-cultures and evidence of NPCs suppression of T cells activated by allogeneic DCs. (A-C) Representative dotplots of data presented in Figure 4; T cell activation displayed as proliferation (x-axis) vs CD25 expression (y-axis) for T cell/NPC co-culture (A), PBMC/NPC co-culture (B) and T cell/moDC/NPC co-culture (C) . (D-F) Untreated and IFNg-stimulated NPCs were added to T cell-alloDC co-cultures to examine suppressive capacity. Representative (B) dotplots, (E) histograms and (C) quantification of proliferation index (PI) and CD25 expression in CD4+ and CD8+ T cells. Red = CD4+ T cells, blue = CD8+ T cells. N=8. Data are represented as mean ± STDEV. One-way ANOVA; n.s. = not significant \*p<0.05, \*\*p<0.01, \*\*\*p<0.005, \*\*\*\*p<0.001. (G) Heatmap of combined data presented in Figure 4, Figure 5 and Sup Figure 6 with relative concentration of human immune cytokine panel relative to untreated TC in T cell/NPC co-culture.

Supplemental Figure 6. The mechanism of suppression was not determined, but several mechanisms common to hPSC were excluded. The contact-dependent (A-G) and -independent (H-N) mechanisms of suppression of T cells by NPCs and IFN-NPCs was investigated. (A) Expression of co-inhibitory molecules in NPCs and IFN-NPCs was measured in the nCounter dataset. (B-C) Blocking the highly expressed molecules, CD276 (B) and TIGIT (C; receptor for CD112, CD113, CD155) with 10 uM blocking antibodies in the 5-day T cell/NPC co-culture had no effect on suppression. (E-F) High gene (E) and surface (F) expression of CD47 was confirmed by qPCR and flow cytometry, respectively, however 10 uM aCD47 blocking antibody had no impact on suppression. (H-J) Expression of immunoregulatory cytokines in NPCs and IFN-NPCs was investigated and TGFb found to be highly expressed. (I) Proof of concept experiment in which 0.1-10 ug/ml rhTGFb successfully suppressed proliferation and CD25 expression in polyclonally stimulated T cells in a dose-dependent manner, however neither blocking of TGFbR1 (using 10 ug/ml SB431542) nor TGFbR1/2 (using 10 ug/ml LY2109761) impacted on the suppressive capacity of NPCs. (K) Expression of IDO-1 and TDO enzymes was measured in the nCounter dataset, with high expression of IDO-1 in IFN-NPCs. (L) However addition of 200 uM of the antagonist 1-L-methyl-tryptophan (1-LMT) or inactive enantiomer 1-D-methyl-tryptophan (1-DMT) had no effect on suppression. (M) Surface expression of CD39 and CD73 on NPCs, compared to hESC and DAn. (N) Blocking of the adenosine pathway via an A2A receptor inhibitor (using 2.5 uM ZM241385) had no effect on suppression. All experiments N=3. Data are represented as mean ± STDEV. One-way ANOVA; n.s. = not significant \*p<0.05, \*\*p<0.01, \*\*\*p<0.005, \*\*\*\*p<0.001.

Supplemental Figure 7. ESC and DAn are neither immunogenic nor immunosuppressive in vitro. (A-C) Representative histograms and quantification of T cell activation represented by proliferation and CD25 expression; untreated and IFNg-stimulated DAn (A-B) and hESCs (C-D) were tested in the T cell co-culture (A/C; n=6) and T cell/moDC co-culture (B/D; n=3). Quantification of proliferation index (PI), using Modfit modelling software, and CD25 expression in CD4+ and CD8+ T cells from all assays. (D) Supernatants from the DAn co-cultures with T cell and T cell/moDC or DAn/IFN-DAn only wells were collected and subjected to the 8-plex human cytokine Luminex panel, measuring the concentration of IFNg, TNFa, IL-2,IL-6, IL-8, IL-4, IL-10, GM-CSF. Heatmap displays cytokine concentration relative to the untreated T cell well. N=3. Data are represented as mean ± STDEV. One-way ANOVA; n.s. = not significant \*p<0.05, \*\*p<0.01, \*\*\*p<0.005, \*\*\*\*p<0.001.
