## Supplementary figures and images for "The immunological profile of RC17 hESC-derived dopaminergic neural progenitor cells *in vitro*: implications for the STEM-PD clinical trial"

### image1.jpg

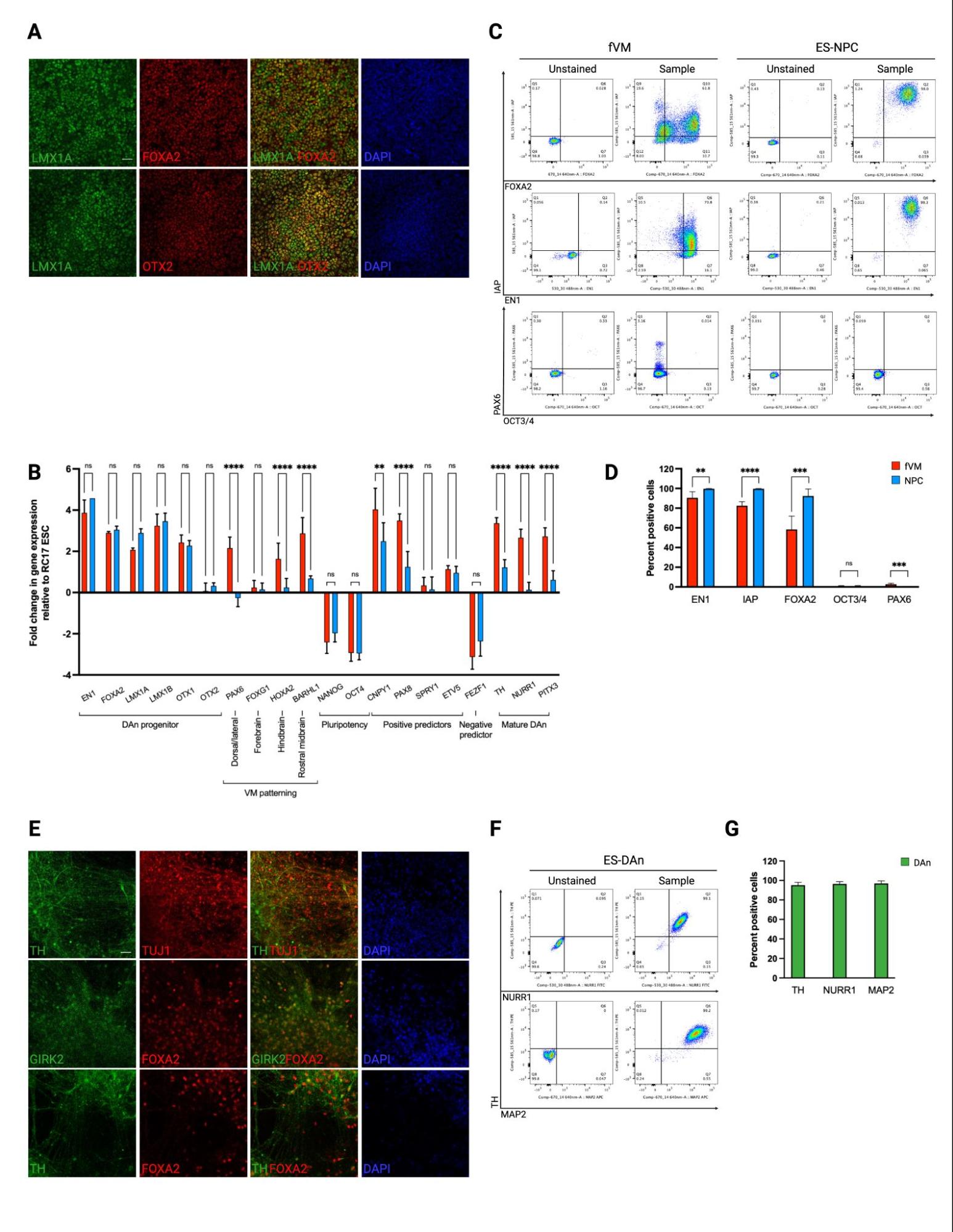

### image2.jpg

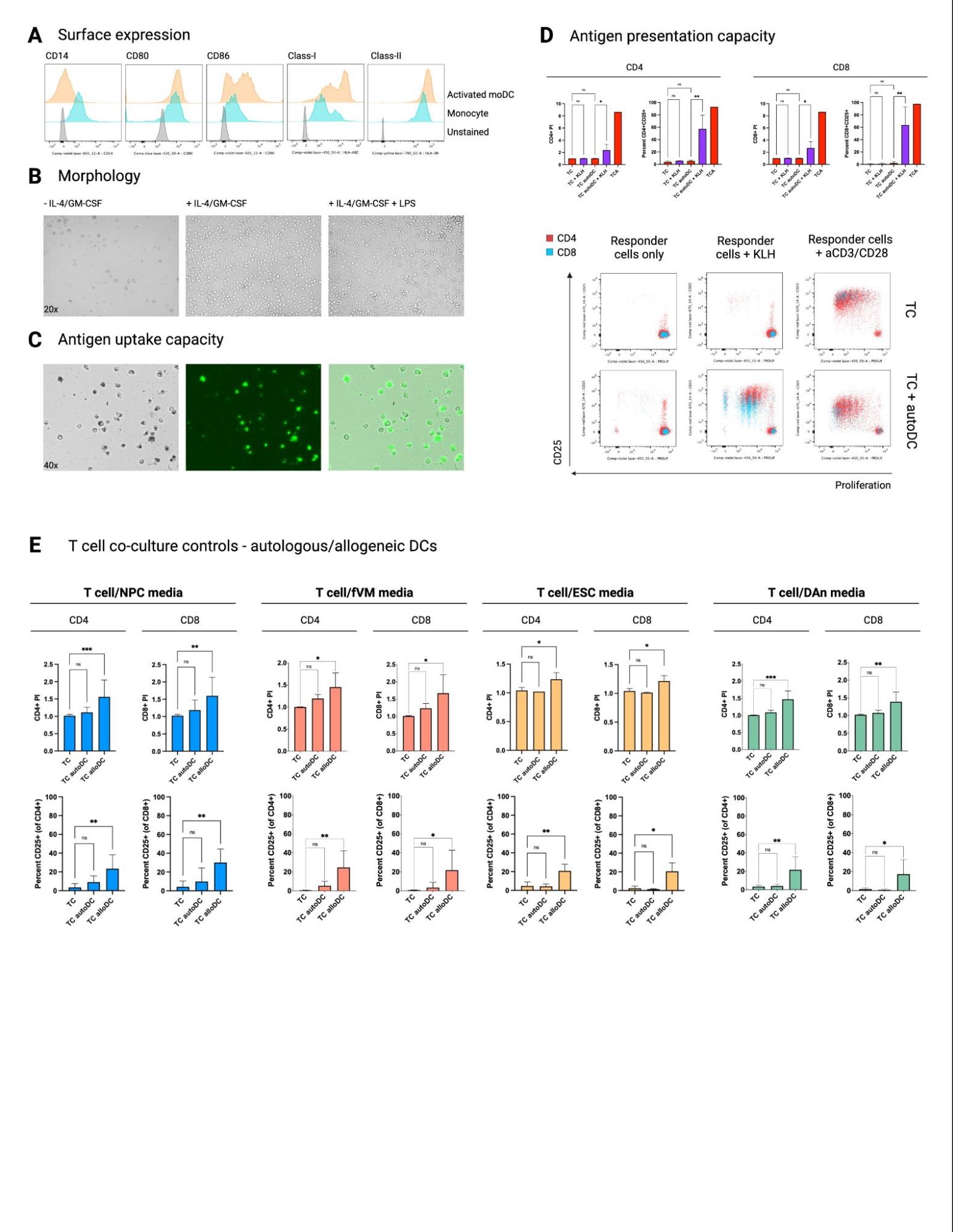

### image3.jpg

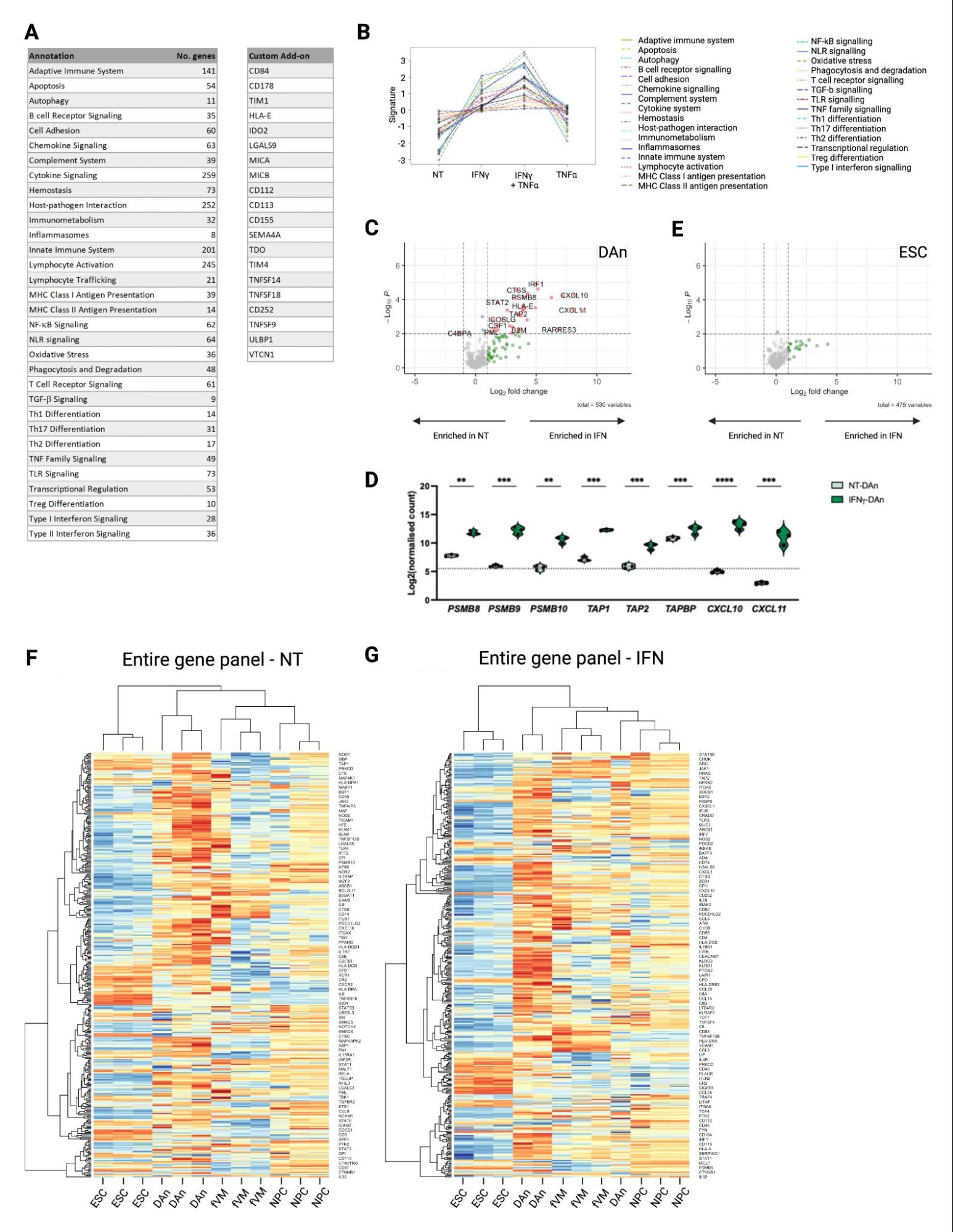

### image4.jpg

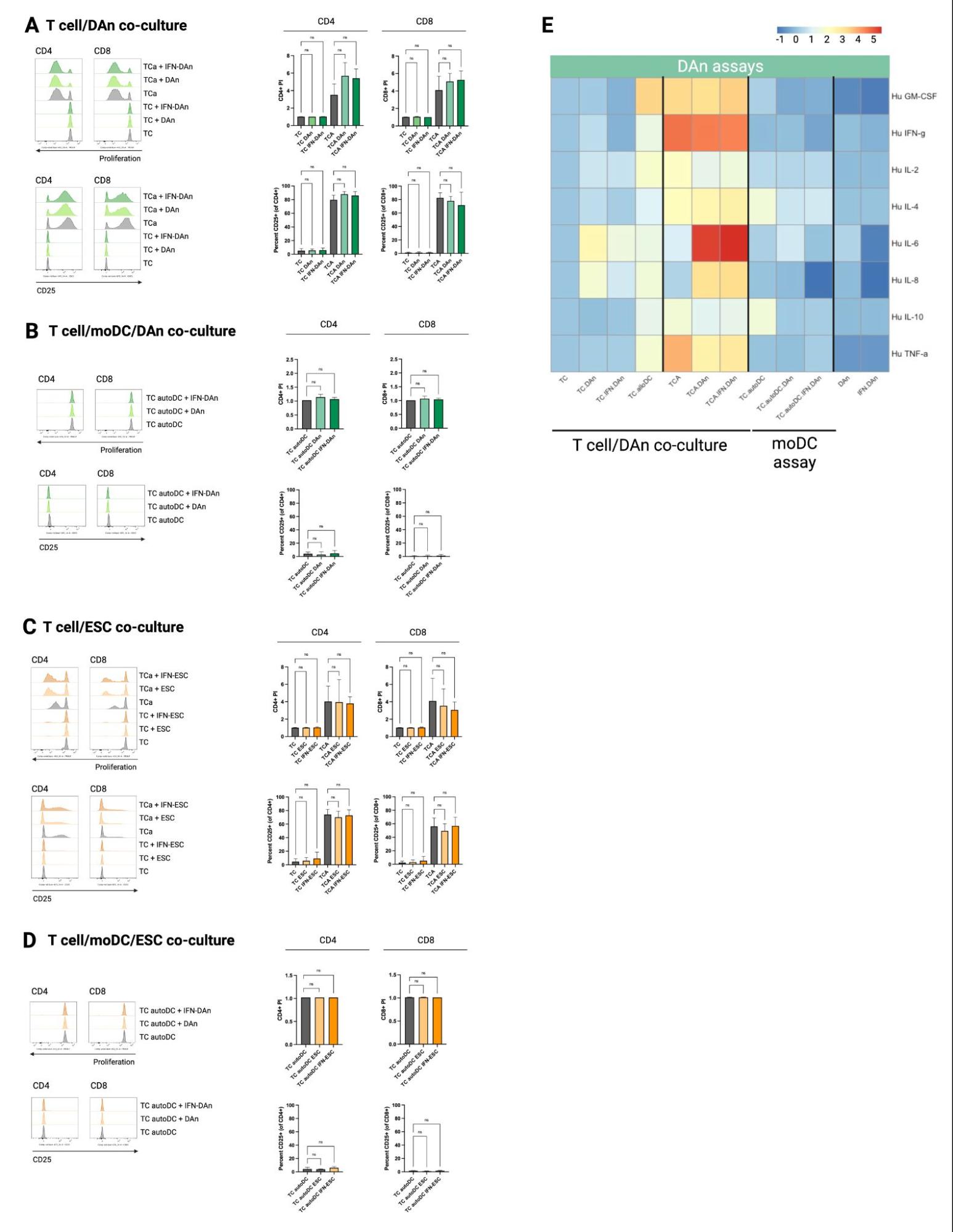

### image5.jpg

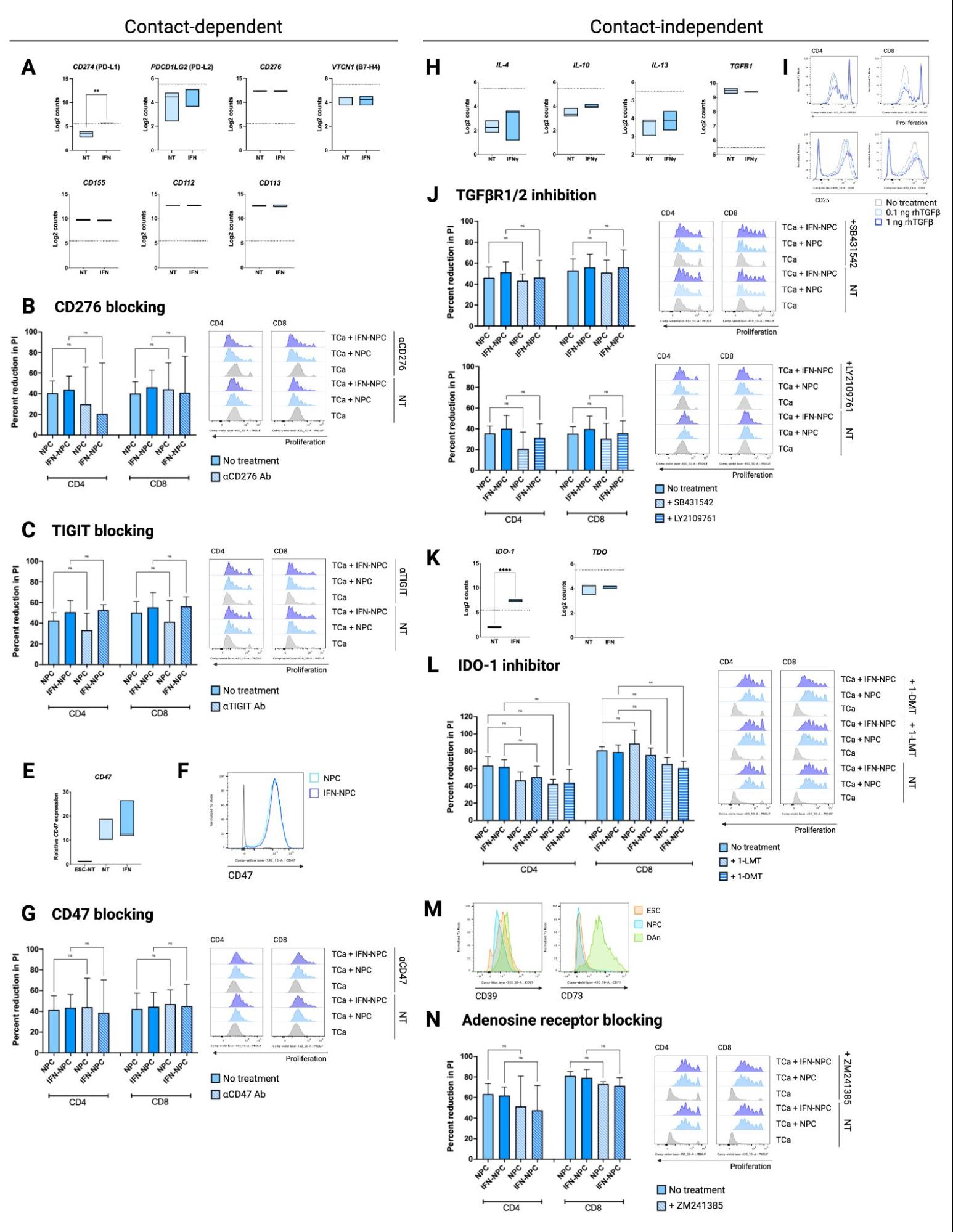

### image6.jpg

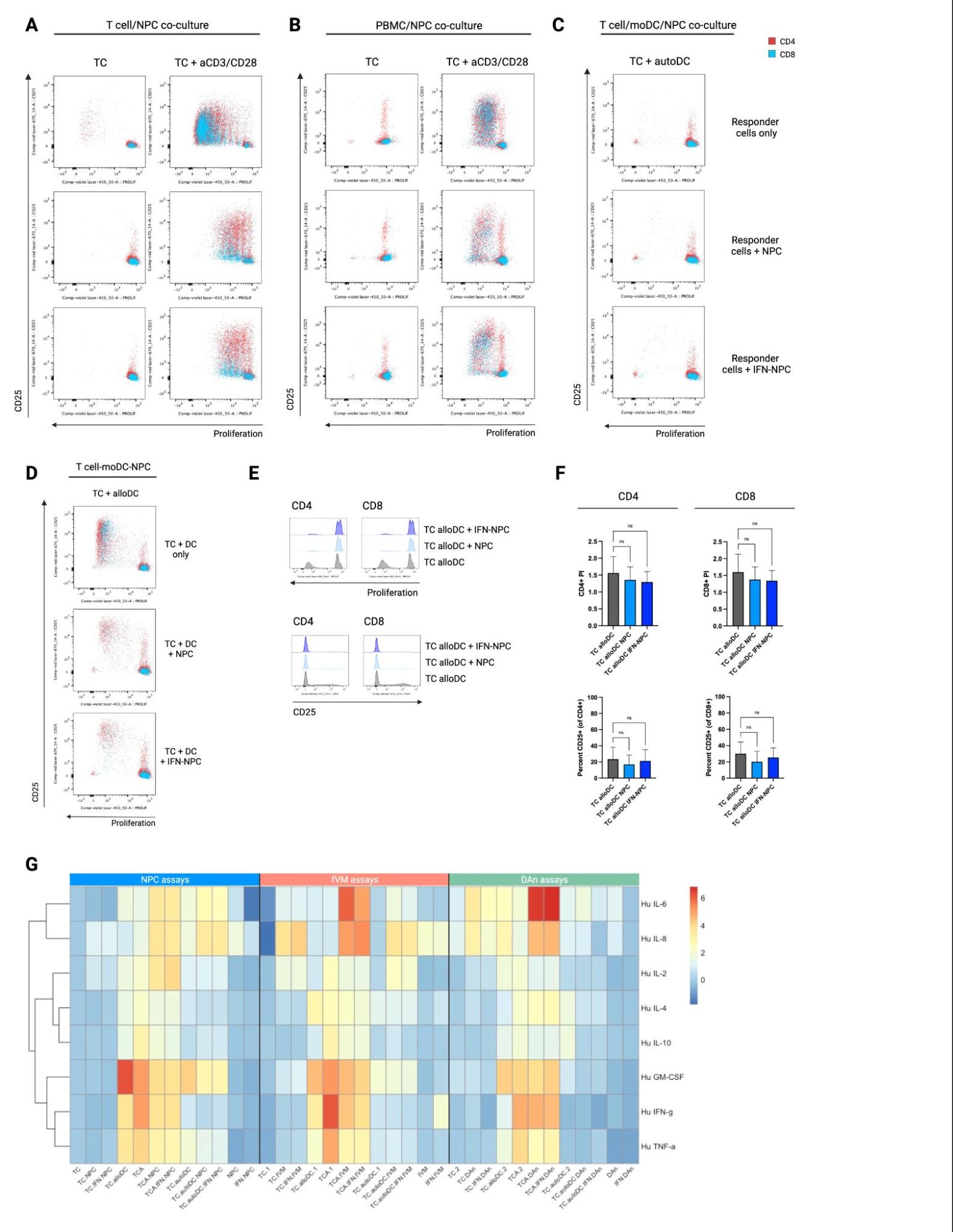

### image7.jpg

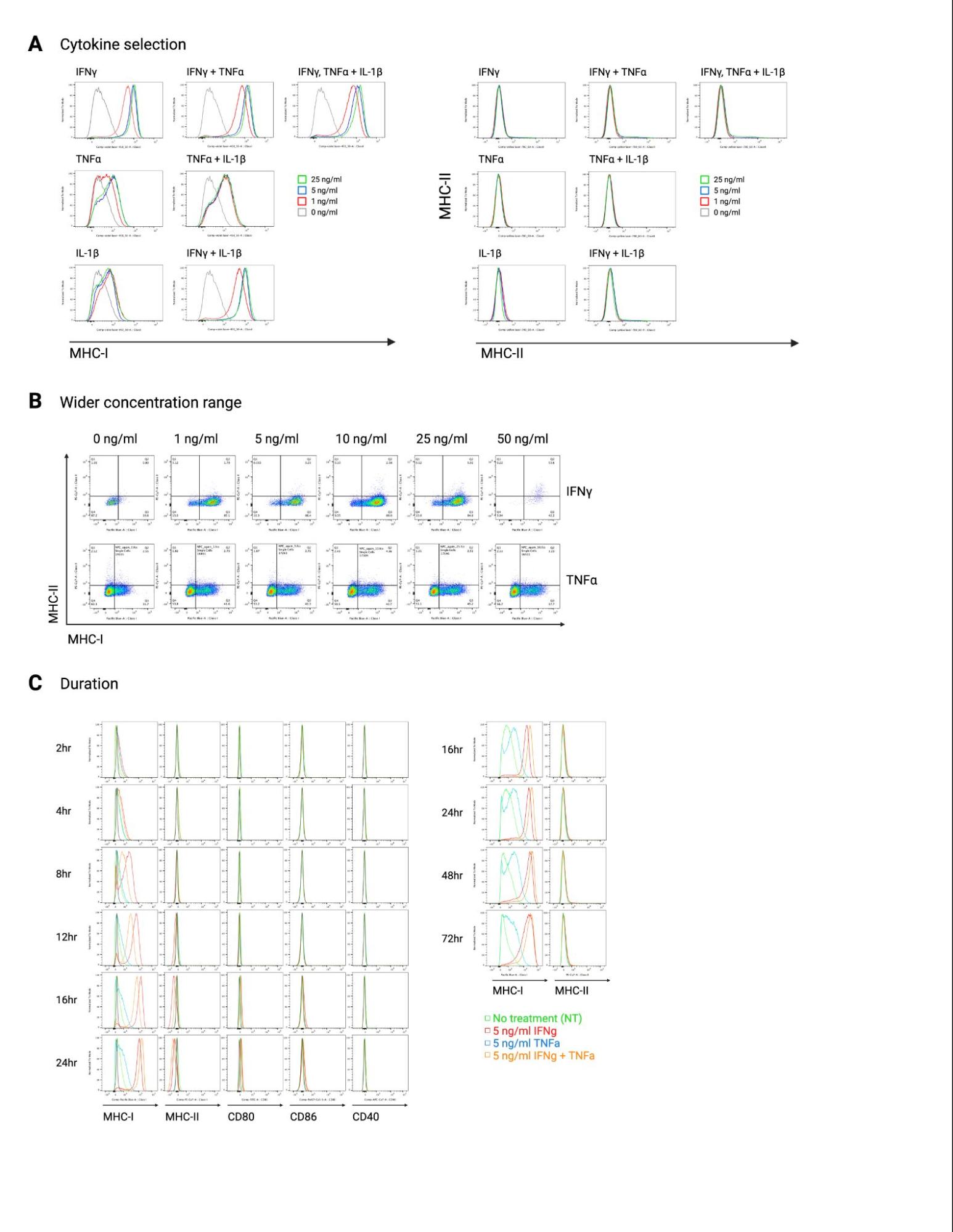
